# RGAST: A Relational Graph Attention Network for Multi-Scale Cell-Cell Communication Inference from Spatial Transcriptomics

**DOI:** 10.1101/2024.08.09.607420

**Authors:** Yuqiao Gong, Zhangsheng Yu

## Abstract

Cell-cell communication (CCC) plays a fundamental role in tissue organization and function. Recent advances in spatial transcriptomics (ST) technologies have enabled high-resolution mapping of CCC at single-cell level. However, existing computational approaches for CCC inference face several limitations, including reliance on predefined ligand-receptor databases, loss of single-cell resolution, and inability to model long range communications. To address these challenges, we developed RGAST, a deep learning framework that integrates both spatial proximity and transcriptional profiles to reconstruct multi-scale CCC networks de novo. In our analysis, RGAST revealed directional communication from peripheral to central nuclei in mouse hypothalamus and tumor invasion signaling axis in breast cancer. Leveraging a relational graph attention network, RGAST effectively captures both local and global communication patterns while learning low-dimensional representations of ST data, which are versatile in multiple downstream tasks. Our results demonstrate that RGAST enhances spatial domain identification accuracy by approximately 10% compared to the second method in 10X Visium DLPFC dataset. Furthermore, RGAST facilitates the discovery of spatially variable genes, enables more precise cell trajectory inference and reveals intricate 3D spatial patterns across multiple sections of ST data.

## Introduction

Spatial transcriptomics (ST) is a breakthrough technology that allows researchers to investigate gene expression and tissue architecture simultaneously in a spatially resolved manner[1]. Technologies such as 10x Visium[2], Slide-seq[3], Stereo-seq[4], MERFISH[5], seqFISH+[6] and HDST[7] enable profiling of gene expression in captured locations, or "spots" at the level of several cells, single cell or even subcellular structures. These breakthroughs have catalyzed the development of computational tools to infer cell-cell communication (CCC), a cornerstone of tissue homeostasis and disease progression. Traditional methods like CellChat [8], NicheNet [9], and CellPhoneDB [10] rely heavily on predefined ligand-receptor databases to infer CCC and suffer from critical drawback: loss of single-cell resolution. While emerging ST-aware tools such as SpaTalk [11] and SpatialDM [12] attempt to leverage spatial proximity, they remain constrained by simplistic distance thresholds and fail to account for distal communication mediated by cytokines or extracellular vesicles.

To address these gaps, we present RGAST (Relational Graph Attention Network for Spatial Transcriptomics), a deep learning framework that integrates spatial proximity and transcriptional profiles to reconstruct multi-scale CCC networks. RGAST employs a hierarchical graph attention mechanism to model both local and global communication patterns, dynamically weighting interactions based on spatial distance and gene expression similarity. Unlike database-dependent methods, RGAST infers CCC networks de novo by adaptively learning connection weight between cells in ST data, enabling the discovery of context-specific signaling axes without prior knowledge of ligand-receptor pairs. Through applications to mouse hypothalamic nuclei and breast cancer microenvironments, we demonstrate RGAST’s ability to uncover directional signaling hubs and tumor-stroma crosstalk, bridging the gap between spatial transcriptomics and functional CCC mapping.

Beyond CCC inference, RGAST’s unified architecture supports versatile downstream analyses. The locally and globally award representations enhance spatial domain identification by capturing spatially coherent transcriptional states, outperforming existing methods by ∼10% in clustering accuracy on 10X Visium DLPFC data. Additionally, RGAST identifies spatially variable genes (SVGs) critical for tissue patterning and facilitates trajectory inference by resolving pseudotemporal ordering of cells within spatially constrained niches. By aggregating interactions across serial tissue sections, RGAST further reconstructs 3D spatial patterns, offering insights into spatially layered tissue structure.

## Results

### Overview of the RGAST model

The spatial distribution of transcriptional expression in tissues is critical for elucidating biological functions and characterizing interactive biological networks [13]. Spatial transcriptomics (ST) data provides valuable location-specific information that can be leveraged effectively. Notably, certain tissue spots, despite being physically distant, may exhibit long-range communication and similar expression patterns due to their participation in spatially regulated processes such as diffusion or signaling [1]. In the RGAST model, we construct a heterogeneous graph by considering two distinct relationships (Methods). The first is based on gene expression similarity between spots, while the second is determined by spatial proximity, accounting for their relative physical locations (Fig. 1). By integrating both relationships, we employ a relation-aware attention mechanism that adaptively learns edge weights between spots, dynamically determining the importance of neighboring spots relative to a target spot. These learned weights reflect the communication potential between two cells/spots and are used to update the target spot’s representation through weighted aggregation of information from connected spots.

**Fig. 1.**
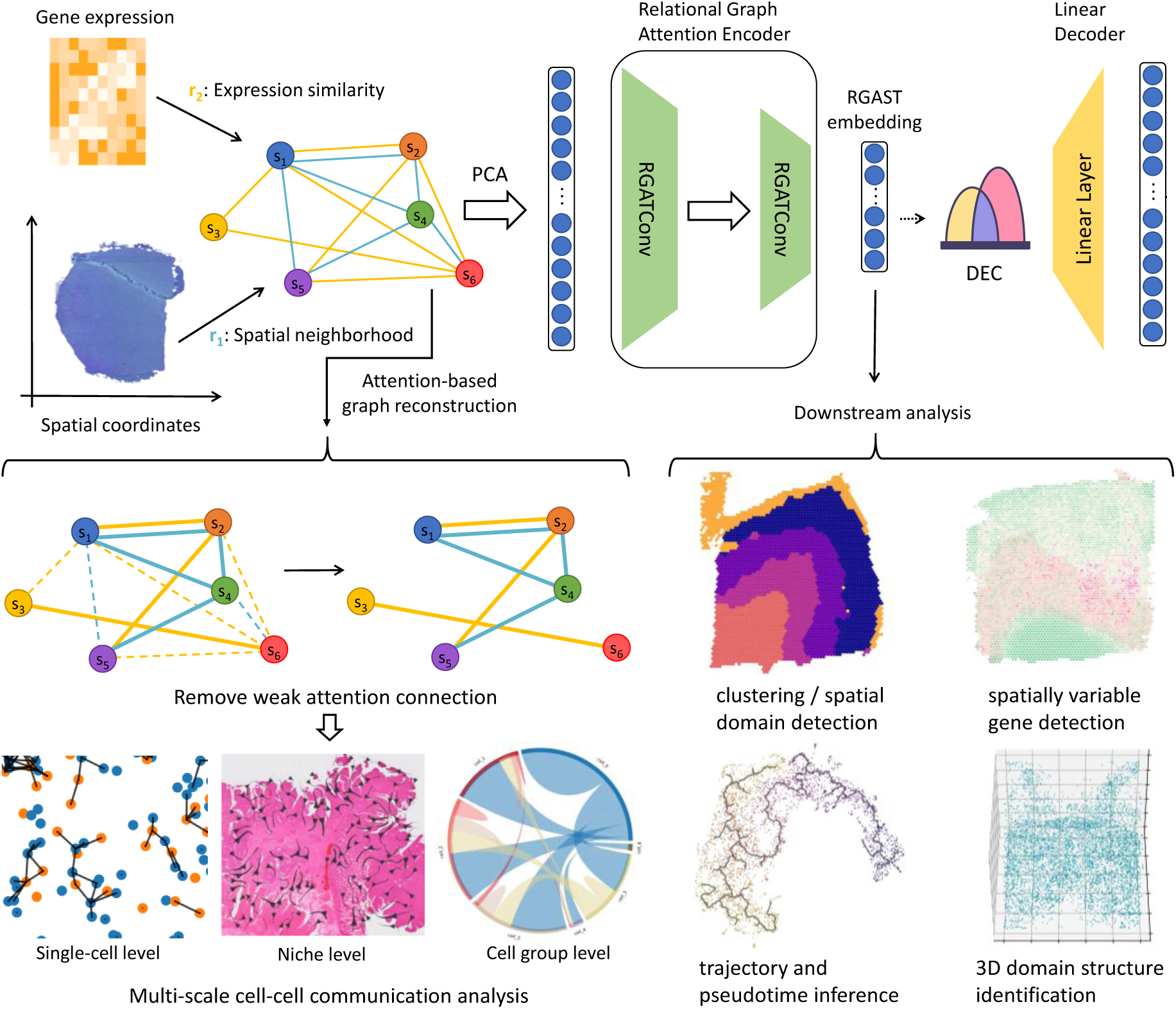
Overview of the RGAST model. RGAST constructs a heterogenous graph using both spatial information and gene expressions. The expression after dimensionality reduction by PCA of each spot is first transformed into a d-dimensional latent embedding by a relational graph attention encoder and then reversed back into a reconstructed expression profile via a linear decoder. An optional unsupervised deep clustering method is then employed to enhance the compactness of the learned latent representation. After training, the graph can be reconstructed by removing edges with low attention score. The reconstructed graph is then used to conduct multi-scale cell-cell communication analysis. The latent embeddings can be applied to many downstream analysis like identifying spatial domains, SVG detection, trajectory inference, and extracting 3D spatial domains.

Using a relational graph attention auto-encoder, RGAST learns cell-cell communication (CCC) pattern and low-dimensional latent embeddings of each spot that encode both spatial and gene expression information (Fig. 1). First, the expression profiles of each spot undergo dimensionality reduction via PCA. An encoder then maps the reduced expression into a d-dimensional latent embedding, which is reconstructed into an expression profile using a linear decoder (Methods). Optionally, an unsupervised deep clustering method can be applied to improve the compactness of the learned latent representation [14] (Methods). Upon convergence, the learned attention scores enable the reconstruction of a CCC graph by filtering out low-confidence edges. The resulting CCC atlas can be aggregated to perform niche-level and cell-type-level CCC analysis (Methods). Furthermore, the derived latent embeddings can enhance various downstream analyses, including data visualization, spatial domain identification, trajectory inference, spatially variable gene (SVG) detection, and 3D spatial domain extraction.

### RGAST uncovers spatially hierarchical signaling in the mouse hypothalamic preoptic region

To assess RGAST’s capability in deciphering cell-cell communication (CCC) at single-cell resolution, we applied it to a mouse hypothalamic preoptic MERFISH dataset comprising 6 tissue slices [15]. For benchmarking, we compared RGAST against several state-of-the-art (SOTA) methods specifically designed for single-cell CCC inference: DeepLinc [16], Scriabin [17], NICHES [18], and COMMOT [19]. Given the absence of a definitive ground truth CCC network, we employed a consensus network as a proxy reference (see Methods). Our results demonstrate that RGAST consistently achieved significantly higher precision across all 6 MERFISH slices (Fig. 2a). Although a modest reduction in recall was observed, the F1 score— which balances both precision and recall—highlighted RGAST’s superior performance (Fig. 2a). This underscores RGAST’s ability to minimize false-positive interactions while preserving biologically meaningful signals. Among the benchmarked methods, RGAST, Scriabin, NICHES, and COMMOT maintained near-perfect accuracy, whereas DeepLinc exhibited comparatively lower performance. This discrepancy can be largely attributed to DeepLinc’s over-sensitivity add-on paradigm in graph reconstructions, which led to an inflated detection of spurious cell connections.

**Fig. 2.**
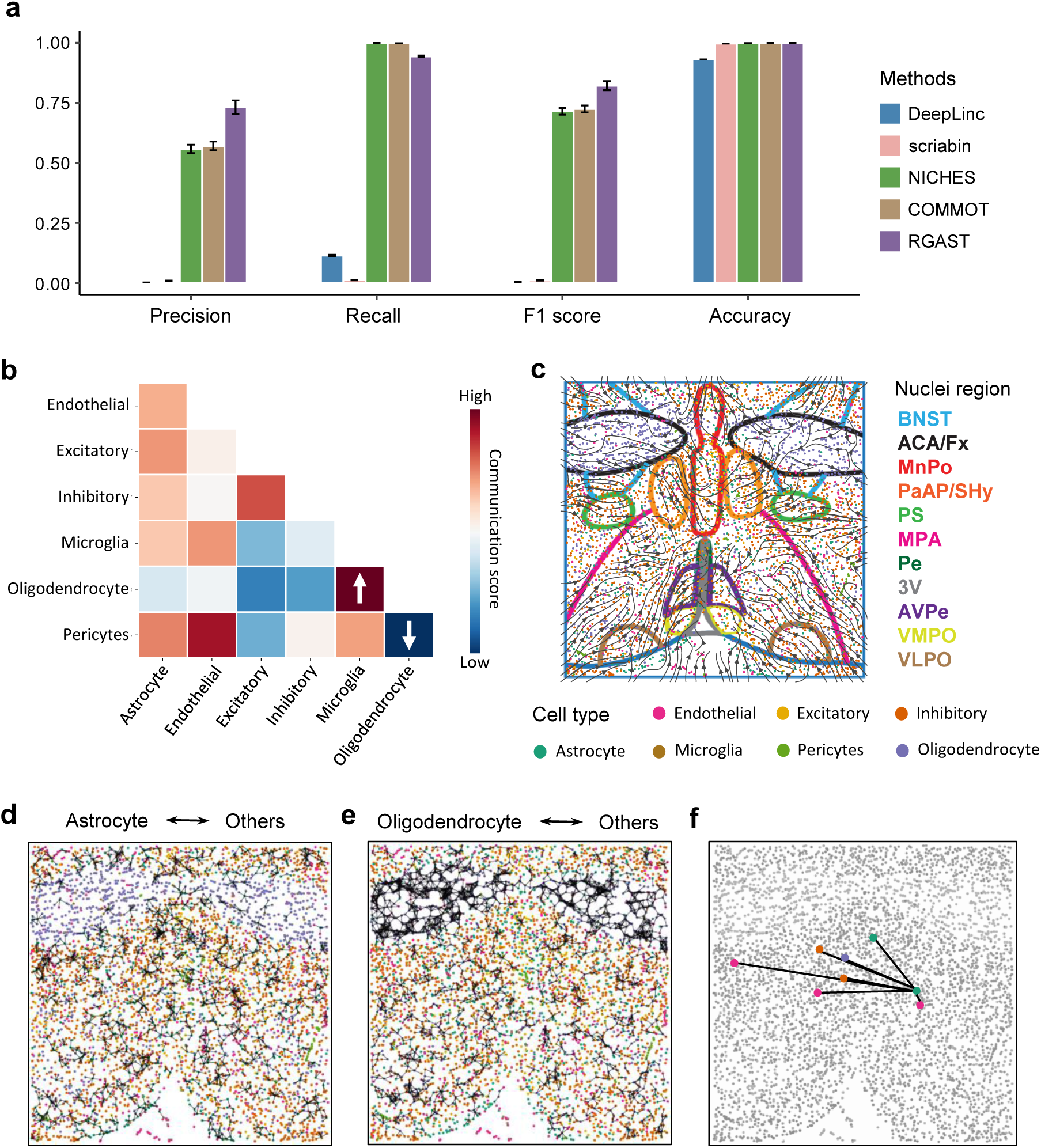
Multi-scale CCC analysis in the mouse hypothalamic preoptic region. **a,** Performance of different methods in constructing single-cell communications network. We run each method on all 6 slices of the MERFISH data. The bar represents mean value, and the error bars represent the standard error calculated across these slices. **b,** Cell type level CCC analysis result on the Bregma +0.26 slice. The up-headed arrow indicates highest communication score, while down-headed arrow indicates lowest communication score. **c,** Niche level communication flow plot of the Bregma +0.26 slice. The cell type annotation and hypothalamic nuclei segmentation are adopted from Moffitt et al. and are colored identically to the nuclei abbreviations listed to the right. BNST, bed nucleus of the stria terminalis; ACA, anterior commissure; Fx, fornix; MnPO, median preoptic nucleus; PaAP, paraventricular hypothalamic nucleus, anterior parvicellular; SHy, septohypothalamic nucleus; PS, parastrial nucleus; MPA, medial preoptic area; Pe, periventricular hypothalamic nucleus; 3V, third ventricle periventricular nucleus; AvPe, anteroventral; VMPO, ventromedial preoptic nucleus; VLPO, ventrolateral preoptic nucleus. **d,** Short-range single-cell resolution communication comprising astrocyte. **e,** Short-range single-cell resolution communication comprising oligodendrocyte. **d,** Long-range single-cell resolution communication between an astrocyte and other cells.

We next conducted a detailed multi-scale cell-cell communication (CCC) analysis on the Bregma +0.26 slice. At the cell-type level, oligodendrocyte-microglia interactions were the most prominent (Fig. 2b), aligning with their well-documented roles in myelin maintenance and neuroinflammatory responses [20–22]. For example, microglia-oligodendrocyte crosstalk via TGFβ1-TGFβR1 is known to mediate myelin compaction [22]. In contrast, oligodendrocyte-pericyte interactions were notably sparse (Fig. 2b), likely due to their spatial segregation—pericytes are primarily localized to vasculature, whereas oligodendrocytes are concentrated in white matter tracts [23, 24]. Astrocytes emerged as central hubs in the CCC network, participating in the majority of interactions (Fig. 2b). This pan-cellular connectivity corroborates astrocytes’ function as metabolic and immune integrators [25]. This observation was further supported at single-cell resolution: astrocytes exhibited widespread spatial distribution, enabling frequent short-range interactions with diverse cell types (Fig. 2d), whereas oligodendrocytes were confined to specific niches and primarily engaged in homotypic interactions (Fig. 2e).

Niche level analysis further uncovered a directional signaling from peripheral hypothalamic nuclei (BNST, PS, MPA, VLPO) to central regions (MnPO, 3V, VMPO) (Fig. 2c). This indicate that MnPO and VMPO, as the central nuclei of preoptic area (POA), may be the core hub for integrating multiple physiological and behavioral signals. For example, median preoptic nucleus (MnPO) coordinates thermoregulation, sleep-wake cycles, and fluid balance [26–28], while the ventromedial preoptic nucleus (VMPO) mediates social behavior and sex hormone signaling [15, 29]. Limbic inputs from regions like BNST and MPA likely convey emotional, stress-related, and environmental information to these hubs for multimodal integration [30]. This organizational framework is supported by recent single-cell transcriptomic evidence showing that POA neurons establish functional specialization during embryonic development, with inter-regional communication networks maturing postnatally through progressive refinement of signaling pathways [31].

Notably, RGAST successfully identified long-range cellular communications that transcend spatial proximity constraints. Our analysis revealed biologically plausible long-range interactions between astrocytes and multiple cell types, including oligodendrocytes, endothelial cells, and inhibitory neurons (Fig. 2f). These findings align with established neurobiological mechanisms: (1) Astrocytes support distal oligodendrocyte function through lactate secretion and BDNF-mediated trophic support, facilitating myelination and axonal metabolic maintenance [32]; (2) Through specialized end-feet structures enveloping vasculature, astrocytes engage in blood-brain barrier formation with endothelial cells while dynamically regulating vascular permeability and cerebral blood flow via VEGF and angiopoietin-1 signaling [33]; (3) Astrocytes maintain neuronal energy homeostasis through long-distance metabolite transport, particularly lactate shuttling [34]. These results demonstrate RGAST’s unique capability to detect spatially extended yet biologically meaningful interactions without imposing molecular preconceptions. The identification of these known neuroglial interaction patterns not only validates RGAST’s spatial resolution but also confirms its ability to reconstruct functional hierarchies within complex neural circuits.

### Spatiotemporal dynamics of cell-cell communication in breast cancer revealed by RGAST

Breast cancer continues to pose a significant challenge in oncology due to its profound molecular heterogeneity and the intricate cellular interactions within the tumor microenvironment [35]. To further evaluate RGAST’s capabilities, we applied it to a highly multiplexed HDST breast cancer dataset. Compared to the MERFISH dataset, all computational methods exhibited reduced precision, recall, and F1 scores, underscoring the increased difficulty of resolving spatial interactions in highly heterogeneous tumor tissues (Fig. 3a). Despite these challenges, RGAST consistently outperformed competing methods in precision and achieved the highest F1 score. Its accuracy, along with NICHES and COMMOT, approached 1 as before (Fig. 3a), demonstrating its robustness in deciphering complex cellular architectures within breast cancer.

**Fig. 3.**
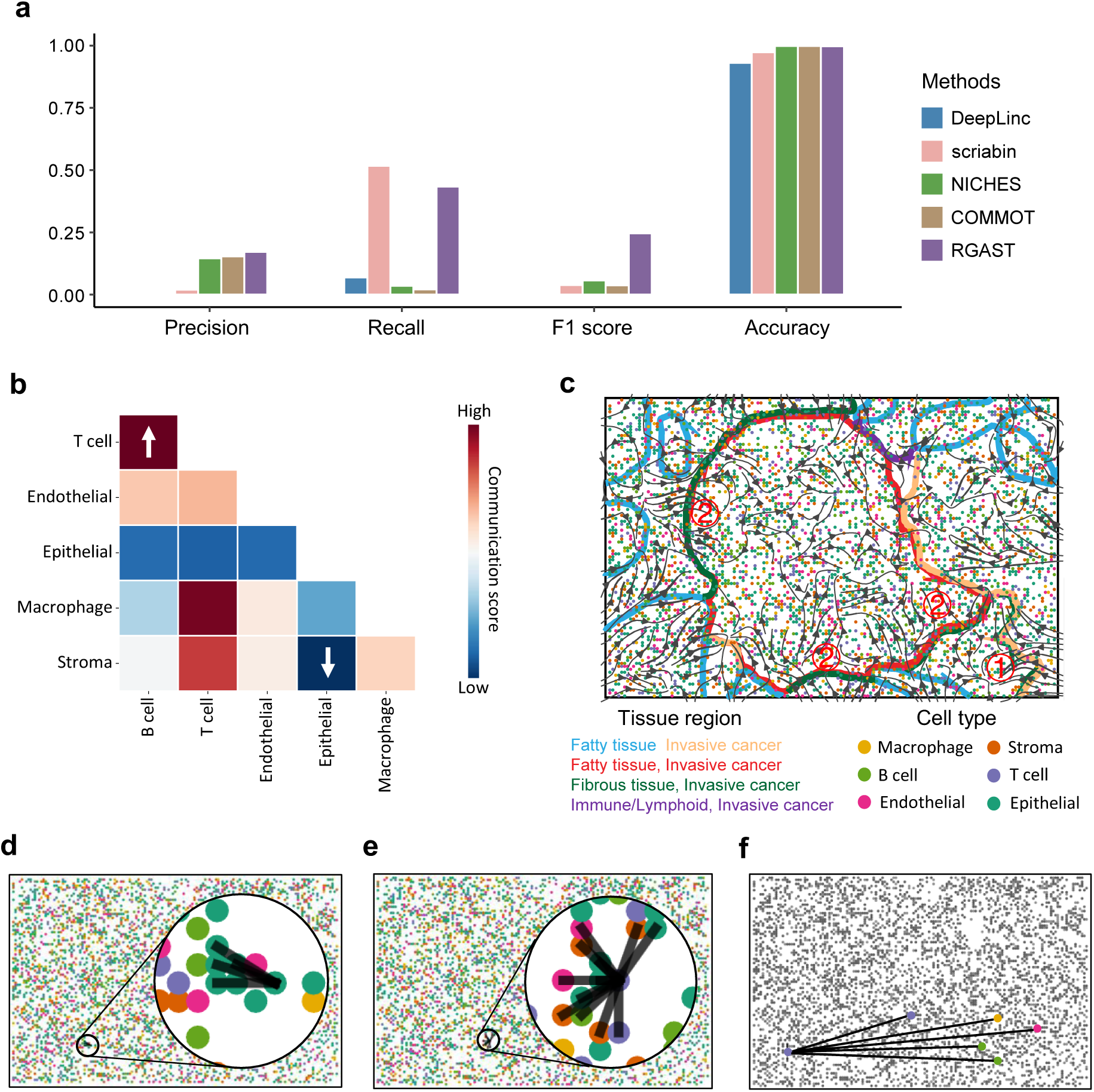
Multi-scale CCC analysis in the HDST breast cancer data. **a,** Performance of different methods in constructing single-cell communications network. **b,** Cell type level CCC analysis result. The up-headed arrow indicates highest communication score, while down-headed arrow indicates lowest communication score. **c,** Niche level communication flow plot. The cell type annotation and tissue region segmentation are adopted from Vickovic et al. and are colored identically to the legend. ① indicates the flow from invasive cancer area to fibrous tissue. ② indicates flow from fibrous tissue to fatty tissue-enriched tumor regions. **d,** Short-range single-cell resolution communication of an epithelial cell. **e,** Short-range single-cell resolution communication of a T cell. **d,** Long-range single-cell resolution communication between a T cell and other cells.

Cell type-level CCC analysis demonstrated that T cells exhibited the strongest interaction scores with B cells, consistent with well-established mechanisms of T-B cell collaboration in adaptive immunity. Specifically, this interaction likely reflects the role of T follicular helper (Tfh) cells, which secrete cytokines such as IL-21 to regulate B cell proliferation and antibody maturation [36]. In contrast, stromal-epithelial interactions were remarkably weak, supporting prior studies that emphasize the metabolic and spatial segregation between stromal fibroblasts and epithelial tumor cells in breast cancer [37]. Notably, epithelial cells displayed the weakest communication with other cell types (Fig. 3b), a pattern further evident at single-cell resolution, where individual epithelial cell primarily engaged in homotypic interactions (Fig. 3d). This aligned with their propensity for clonal expansion and limited crosstalk with other lineages [38]. Conversely, T cells exhibited strong communication scores with most cell types (Fig. 3b), a trend observable even at single-cell resolution: these cells displayed extensive heterotypic interactions with neighboring cells (Fig. 3e), reinforcing their role as central immune coordinators within the tumor microenvironment (TME) [39].

Our niche-level spatial analysis uncovered a prominent directional signaling axis, extending from the invasive cancer periphery toward the fibrous tissue interface (① in Fig. 3c), followed by subsequent signaling propagation into central fatty tissue-enriched tumor regions (② in Fig. 3c). This spatial progression mirrors the multistep cascade of metastatic dissemination, wherein invasive tumor cells exploit stromal remodeling, metabolic symbiosis, and immune-evasive niches to facilitate distal colonization [40]. At the invasive cancer-fibrous tissue boundary, the observed signaling flux likely reflects CAF-driven ECM remodeling, which generates biomechanical guidance cues for collective tumor cell migration [41]. These fibrous regions—enriched with cross-linked collagen IV and aligned ECM fibers—establish a stiffened microenvironment that promotes tumor cell protrusion formation and directional motility. This aligns with the "pulse mode" metastasis model, wherein cyclic ECM stress accumulation drives tumor nest fragmentation and dissemination [42]. Further signaling convergence into fatty tissue niches may correspond with adipocyte-mediated metabolic reprogramming, as tumor cells upregulate fatty acid oxidation (FAO) pathways to exploit lipid-rich microenvironments for hypoxic stress survival [43]. This spatial progression—from ECM degradation at the invasive front to metabolic adaptation in fatty niches—recapitulates the "seed-soil" hypothesis, highlighting how tumor cells dynamically engage with organ-specific microenvironments to complete metastatic colonization.

Notably, RGAST identified long-range interactions between individual T cells and distal macrophages, endothelial cells, and B cells (Fig. 3f). This aligns with studies demonstrating cytokine-mediated distal communication (e.g., CCL5-CCR5 axis) and exosome trafficking across tumor niches [44, 45]. For instance, tumor-associated macrophages (TAMs) secrete IL-10 to remotely suppress T cell cytotoxicity [46], while endothelial cells guide immune cell trafficking via S1P gradients [47]. These findings underscore the utility of RGAST in capturing both local and systemic communication networks within spatially complex tumors.

### RGAST improves spatial domain detection in various ST technologies with scalability

Beyond cell-cell communication (CCC) analysis enabled by RGAST’s graph reconstruction mechanism, the learned latent embeddings—which integrate both spatial neighborhood relationships and gene expression similarities—demonstrate broad utility across diverse downstream applications. To evaluate RGAST’s spatial clustering performance, we first applied it to spatial transcriptomics (ST) datasets generated using multiple technologies. Our initial benchmark utilized the 10x Visium platform, analyzing spatial expression profiles from 12 human dorsolateral prefrontal cortex (DLPFC) tissue sections[48]. Maynard et al. manually annotated the DLPFC layers and white matter (WM) based on morphological features and gene markers, which we adopted as the ground truth. To compare the clustering accuracy of RGAST with other methods, we employed the adjusted Rand index (ARI) (Methods) and compared it with the non-spatial clustering method Seurat and seven recently developed spatial clustering approaches: SpaGCN[49], BayesSpace[50], SEDR[51], STAGATE[52], conST[53], STMGCN[54] and GraphST[55]. To ensure a fair comparison, we set the maximum number of clusters for each method the same as the ground truth label and used the recommended default parameters for all other methods. For methods that generated latent embedding (i.e., SEDR, STAGATE, conST, GraphST and RGAST), we utilized the Leiden clustering algorithm (Methods). The results demonstrated that RGAST effectively identified the expected cortical layer structures and achieved significant better performance across all slices (Fig. 4b and Supplementary Fig. S1-S11). For instance, in DLPFC section 151675, RGAST clearly delineated the layer borders and achieved the best clustering accuracy (ARI = 0.633). In contrast, the clustering assignment of the non-spatial method Seurat was discontinuous, with many outliers that impeded its clustering accuracy. Not surprisingly, algorithms that leveraged spatial information substantially outperformed the non-spatial clustering method Seurat (Fig. 4b). These findings underscore the superiority of RGAST in spatial domain identification and the necessity to incorporate spatial information in spatial clustering analysis.

**Fig. 4.**
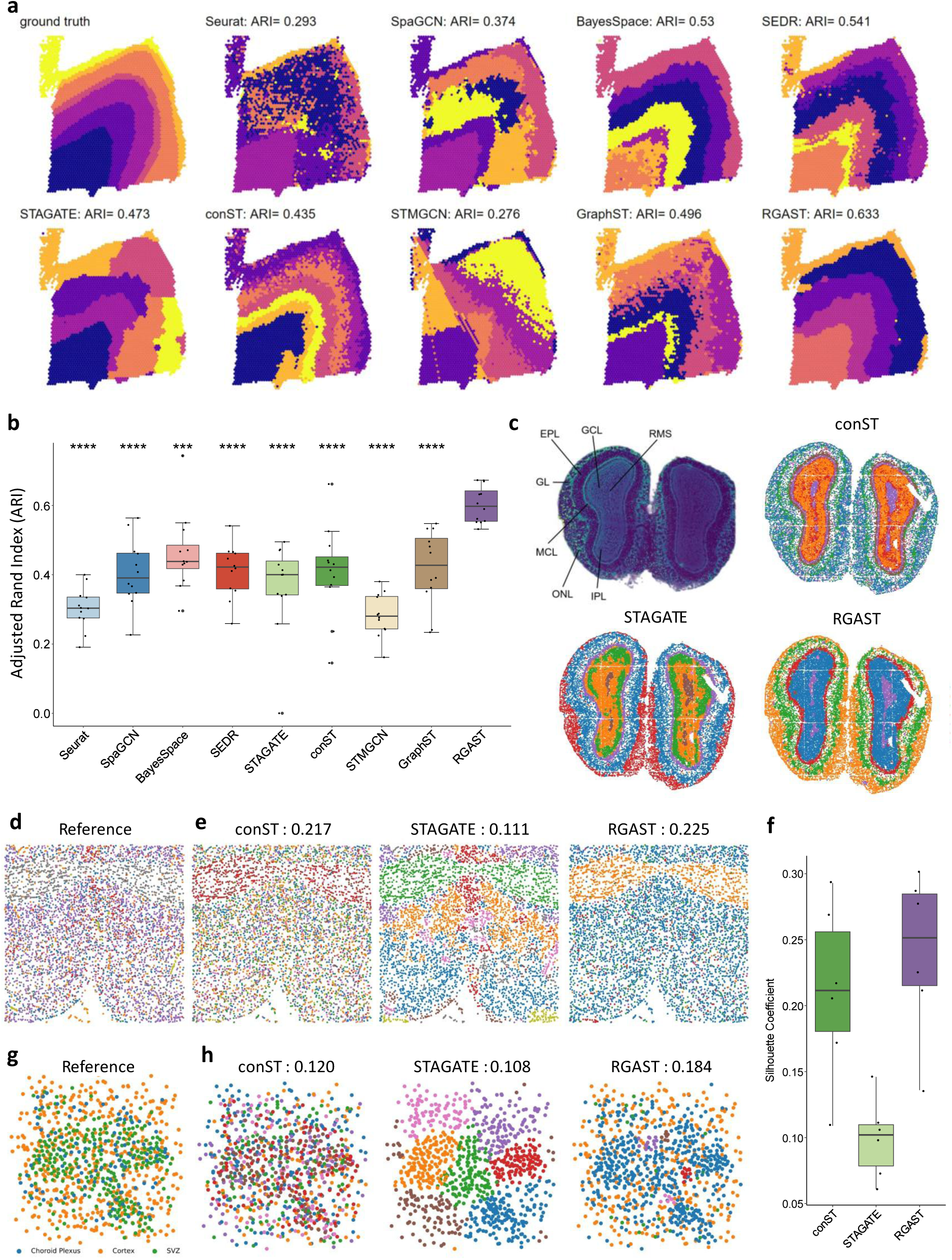
RGAST improves spatial domain detection in various ST technologies. **a,** Clustering results of different methods in the 10X Visium DLPFC section 151675 dataset. **b,** Boxplot of clustering accuracy for 9 methods across all 12 sections of the 10X Visium DLPFC datasets, measured in terms of ARI scores. The reference method (’RGAST’) is compared against other methods using a paired Wilcoxon signed-rank test (one-sided, alternative hypothesis: ARI < RGAST). Benjamini-Hochberg adjusted significance levels are denoted as: (****, p < 0.0001; ***, p < 0.001; **, p < 0.01; *, p < 0.05; ‘ns’, not significant). In the boxplot, the center line, box limits, and whiskers represent the median, upper and lower quartiles, and 1.5× interquartile range, respectively. These parameters are consistent across all boxplots in this paper. **c,** Spatial domains generated by Leiden clustering with a maximum of seven clusters on the low-dimensional conST, STAGATE, and RGAST embeddings in the Stereo-seq mouse olfactory bulb tissue section. Top left panel is the laminar organization of the mouse olfactory bulb annotated in the DAPI-stained image generated by Stereo-seq. RMS, rostral migratory stream; GCL, granule cell layer; IPL, internal plexiform layer; MCL, mitral cell layer; EPL, external plexiform layer; ONL, olfactory nerve layer. **d,** Cell annotation by Moffitt et al. based on scRNA-seq profiling. **e,** Spatial domains generated by different methods in the mouse hypothalamic preoptic MERFISH data of section Bregma +0.26. The number in the title represents the SC score of the different methods, which is consistent with **h**. **f,** Boxplot of clustering accuracy for different methods across all six sections of the MERFISH mouse hypothalamic preoptic datasets, measured in terms of SC scores. **g,** Cell annotation by area, including the Choroid Plexus, subventricular zone (SVZ), and cortex area in the mouse cortex SeqFISH+ dataset. **h,** Spatial domains generated by different methods in the mouse cortex SeqFISH+ dataset.

Next, we compared the performance of RGAST with conST and STAGATE, which have been demonstrated to be superior to other techniques[52, 53], using the Stereo-seq Mouse olfactory bulb dataset. This emerging spatial omics technology can achieve subcellular spatial resolution using DNA nanoball patterned array chips, and the data used here were binned into a cellular-level resolution of approximately 14 μm[4]. Fu et al. annotated the laminar organization of the coronal mouse olfactory bulb in the DAPI-stained image[51](Fig. 4c). Since this dataset contains approximately 20,000 spots, we employed a recently developed Divide-Iterate-Conquer (DIC) strategy to prevent GPU memory exhaustion during training [56], thereby enhancing the scalability of RGAST (Methods). Compared to the clusters identified by conST, those identified using both STAGATE and RGAST embeddings more accurately reflected the laminar organization and corresponded well to the annotated layers. However, STAGATE erroneously separated the GCL into two clusters, whereas RGAST did not (Fig. 4c). Furthermore, the five main layer structures identified by RGAST were all validated by the corresponding marker genes (Supplementary Fig. S12).

We also conducted experiments using Mouse hypothalamic preoptic MERFISH and Mouse cortex seqFISH+ data, which were captured at a higher resolution to achieve single-cell resolution, but no morphology data was available. Since no gold standard was available, we used Silhouette Coefficient (SC) to evaluate the clustering performance of each method. For the MERFISH datasets, Moffitt et al. annotated the cell types based on scRNA-seq expression data[15] (Fig. 4d). The results showed that RGAST consistently outperformed the other methods in all six slices, with STAGATE far behind (Fig. 4e, f and Supplementary Fig. S13-S17). For example, in slice Bregma +0.26, RGAST was more consistent with the clustering results generated by the expression profile (Fig. 4e). In the Mouse cortex seqFISH+ dataset, the spots were annotated as the Choroid Plexus, subventricular zone (SVZ), and cortex area [6] (Fig. 4g). RGAST still demonstrated better performance than the other two methods (Fig. 4h). It’s noteworthy that STAGATE appeared to overuse the spatial neighborhood relationship, as it showed obvious clustering characteristics of block aggregation in such data with discrete cell type distribution (Fig. 4e, h).

### RGAST assist identifying spatially variable genes

Spatially variable genes (SVGs) exhibit varying expression patterns across distinct spatial locations. Identifying SVGs can provide valuable insights into the biological processes that underlie the development and function of tissues and organs. To this end, we incorporated SpaGCN, a tool designed specifically for detecting SVGs, into our benchmarking alongside conST and STAGATE. We compared the performance of these methods to that of RGAST using datasets generated by different ST techniques. In order to ensure a fair and accurate comparison, we utilized identical detection methods and default parameters as those employed by SpaGCN (see Methods section).

We first utilized the section 151674 of DLPFC dataset from 10X Visium data as our benchmarking dataset. Our results demonstrated that SVGs exclusively detected by RGAST exhibited a clear spatial distribution pattern, whereas the spatial distribution patterns of SVGs exclusively detected by other three methods are less discernable (Fig. 5a). We employed Moran’s I and Geary’s C statistics to quantify the degree of spatial autocorrelation of gene expression. A higher Moran’s I statistic and a lower Geary’s C statistic indicate a more distinct spatial pattern. We can see the SVGs identified by RGAST are significantly more reliable than those identified by the other methods (Fig. 5b).

**Fig. 5.**
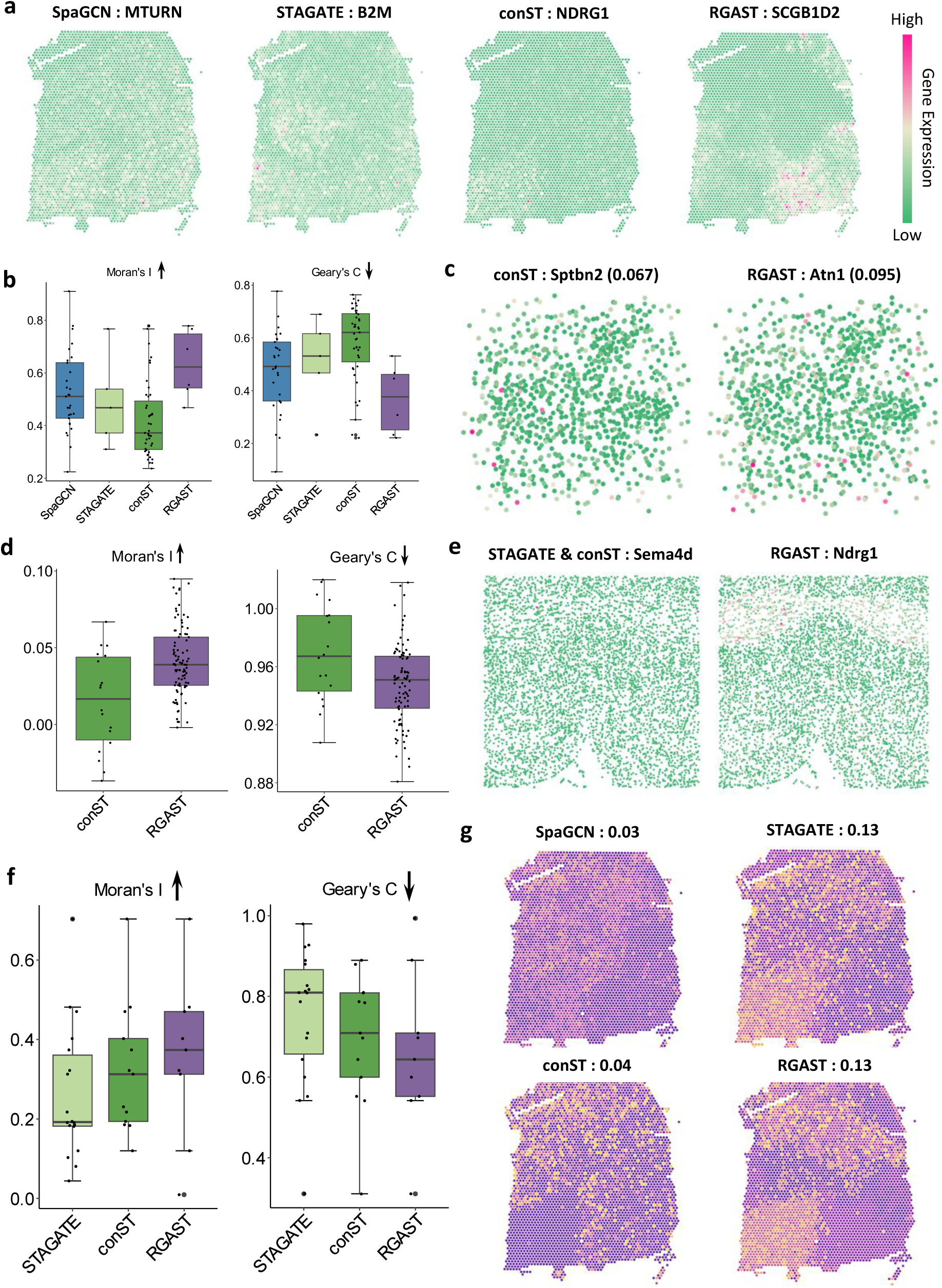
RGAST assist identifying spatially variable genes. **a,** Examples of the spatial distribution of SVGs exclusively detected by SpaGCN, STAGATE, conST and RGAST, respectively, in the DLPFC dataset section 151674. Shades of red indicate the target domain used to detect the corresponding SVGs. Upward arrow means higher value is better, while downward arrow means lower value is better. **b,** Boxplot of Moran’s I and Geary’s C statistics for all SVGs detected by each method in the DLPFC dataset section 151674. **c,** Spatial distribution of SVGs detected by conST and RGAST with the highest Moran’s I in mouse cortex SeqFISH+ data. STAGATE failed to detect any SVGs. **d,** Boxplot of Moran’s I and Geary’s C statistics for all SVGs detected by each method in mouse cortex SeqFISH+ data. **e,** Spatial distribution of SVGs detected by STAGATE conST and RGAST, respectively, in mouse hypothalamic preoptic MERFISH data slice Berman +0.26 with a similar target domain. **f,** Boxplot of Moran’s I and Geary’s C statistics for all SVGs detected by each method in mouse hypothalamic preoptic MERFISH data slice Berman +0.26. **g,** Clustering results using only four SVGs detected by each method as features. The number in the title denotes the ARI score.

Due to the absence of pathological images in the MERFISH and SeqFISH+ datasets, SpaGCN was not included in our follow-up comparison. Interestingly, STAGATE failed to detect any SVGs in the mouse cortex SeqFISH+ dataset, also indicating an unclear spatial domain pattern in this dataset (Fig. 5c). To address this issue, we visualized the SVGs detected by RGAST and conST with the highest Moran’s I, which revealed the superior performance of RGAST (Fig. 5c). In fact, RGAST outperformed conST even when we considered all the SVGs detected by each method (Fig. 5d). For the MERFISH data, we used slice Berman +0.26 as our benchmarking dataset and found that SVGs identified by RGAST displayed a clearer spatial distribution pattern, even when the target domain was similar (Fig. 5e). This was confirmed by our quantitative comparison as well (Fig. 5f). Lastly, to evaluate the representational ability of SVGs detected by each method, we used the SVGs as features to conduct clustering on the DLPFC data, with ARI as our evaluation criterion. For a fair comparison, we randomly selected four SVGs for each method, since STAGATE only detects four SVGs. Our results showed that RGAST and STAGATE performed similarly and significantly better than SpaGCN and conST (Fig. 5g).

### Cell trajectory and 3D spatial model analysis with RGAST

RGAST embeddings, incorporating both local and global awareness, can also be effectively utilized for downstream applications including trajectory inference and 3D spatial clustering analysis. Trajectory inference is a method used to infer the pattern of cells in dynamic developmental processes, with pseudotime representing the progression through this process. Improving the quality of the embedding method can significantly enhance the accuracy and resolution of trajectory inference results[57]. In this study, we selected the DLPFC dataset for trajectory and pseudotime inference, as this tissue represents a natural developmental progression. We used section 151676 as the benchmarking data and compared three methods - SEDR, STAGATE, and conST - with RGAST, as these methods have shown superior performance in trajectory inference analysis[51–53]. To validate the quality of the learned embeddings, we used Monocle3[58] to produce trajectory and pseudotime inference results with embeddings generated from different methods. For pseudotime inference, we selected white matter (WM) as the starting point (Methods).

Based on the results, the UMAP visualization generated by the SEDR embedding appeared as a circular structure in which layer 1 and layer 6 are connected to each other (Fig. 6a). In contrast, the conST and RGAST embeddings showed a streamlined expansion of different layers, while the UMAP visualization of the STAGATE embedding appears clumped together without an obvious linear structure (Fig. 6a). In the inferred development trajectory and pseudotime, STAGATE exhibited too many branching points, and the visualization in the spatial context confirmed the incorrect developmental trajectory inferred by STAGATE (Fig. 6b, c). While the pseudotime inferred by the other three methods reflects the correct "inside-out" developmental ordering of cortical layers, the developmental pseudotime inferred by SEDR and conST is not as smooth as RGAST and exhibits more outliers in each layer structure (Fig. 6b, c). Therefore, our results suggest that RGAST can reveal cell trajectories more accurately than the other methods.

**Fig. 6.**
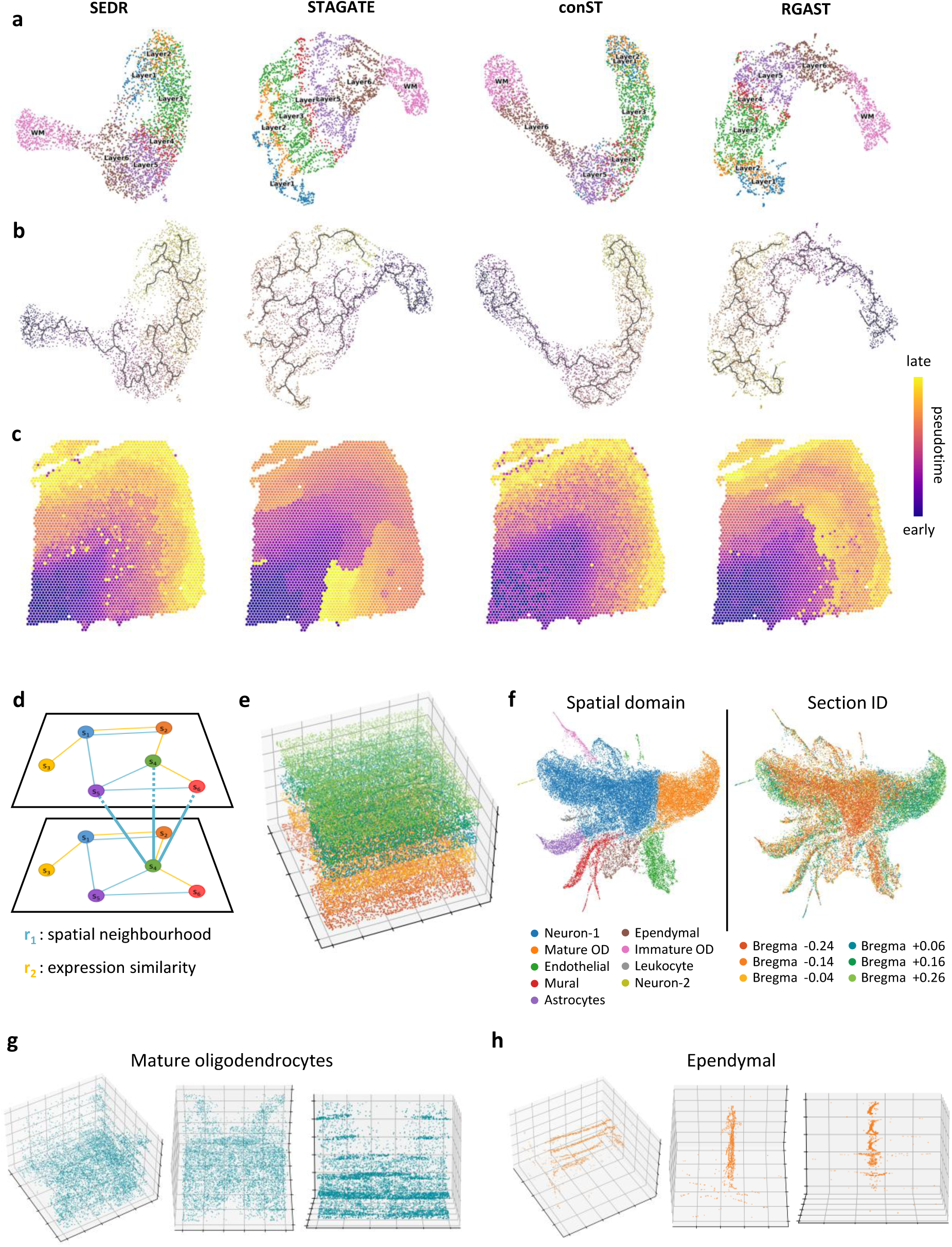
Cell trajectory and 3D spatial model analysis with RGAST. **a,** UMAP visualizations colored by different layers generated by SEDR, STAGATE, conST, and RGAST embeddings, respectively, in the DLPFC section 151676. The order of plots for different methods in b and c is consistent with a. **b,** UMAP visualizations colored by pseudotime inferred from SEDR, STAGATE, conST, and RGAST embeddings, respectively. **c,** Pseudotime inference results from different methods depicted on the spatial physical context. **d,** The 3D RGAST is constructed by by simultaneously considering the 2D relationships within each section and the spatial neighborhoods between adjacent sections. **e,** The aligned six mouse hypothalamic preoptic sections profiled using MERFISH. **f,** Left is UMAP visualization of the RGAST embedding using the 3D model colored by manually annotated spatial domain. Right is UMAP visualization of the RGAST embedding using the 3D model colored by different section. **g,** Visualization of the 3D structure pattern of mature oligodendrocytes from different angles. **h,** Visualization of the 3D structure pattern of Ependymal from different angles.

A single slice of ST data can provide a 2D spatial pattern of a specific organ. When multiple consecutive slices are available for the same organ, we can leverage the relationships between these slices to achieve a more precise representation of each spot. In this study, we extended RGAST to handle 3D ST data by simultaneously considering the 2D relationships within each section and the spatial neighborhoods between adjacent sections (Fig. 6d, Methods). To demonstrate the effectiveness of our approach, we applied RGAST to a pseudo-3D ST dataset obtained by aligning six mouse hypothalamic preoptic sections profiled using MERFISH (Fig. 6e).

Based on the clustering results in the spatial domain, we manually annotated the cell type of each domain using marker genes (Fig. 6f and Supplementary Fig. S18). The UMAP visualization of RGAST embeddings demonstrates effective integration of spots across different tissue sections (Fig. 6f). This integration performance suggests RGAST’s capability to mitigate technical artifacts, including batch effects In the 3D spatial domain results detected by RGAST, a clear structure pattern was observed for mature oligodendrocytes and ependymal cells (Fig. 6g, h), which is consistent with previous studies[15]. Additional visualizations of other spatial domains can be found in Supplementary Fig. S19. These results illustrate that RGAST can facilitate the reconstruction of 3D tissue models and accurately capture 3D expression patterns by incorporating spatial information.

## Discussion

The advent of spatial transcriptomics (ST) has revolutionized our ability to interrogate cellular interactions within their native tissue context. However, existing computational methods for inferring cell-cell communication (CCC) remain constrained by reliance on predefined ligand-receptor databases, limited resolution, and an inability to model long-range interactions. Here, we present RGAST, a deep learning framework that overcomes these limitations by integrating spatial proximity and transcriptional profiles through a relational graph attention mechanism. Our results demonstrate that RGAST not only reconstructs CCC networks with high precision but also enhances downstream analyses, including spatial domain identification, trajectory inference, and 3D tissue modeling.

RGAST’s key innovation lies in its ability to infer CCC networks de novo without prior knowledge of ligand-receptor pairs. Unlike database-dependent tools (e.g., CellChat, NicheNet), which are limited by incomplete interaction annotations, RGAST dynamically learns context-specific signaling patterns by adaptively weighting edges based on gene expression similarity and spatial distance. This approach proved particularly effective in identifying long-range interactions, such as astrocyte-oligodendrocyte crosstalk in the hypothalamus and T cell-macrophage communication in breast cancer, which cannot be identified by locally-constraint methods. The superior performance of RGAST in benchmarking (higher F1 scores vs. DeepLinc, Scriabin, and COMMOT) underscores its robustness in capturing both local and systemic communication networks. Application of RGAST to the mouse hypothalamus uncovered a directional signaling hierarchy from peripheral nuclei (BNST, MPA) to central hubs (MnPO, VMPO), aligning with their roles in integrating physiological and behavioral signals. In breast cancer, RGAST revealed a metastatic signaling axis from invasive fronts to lipid-rich niches, recapitulating the "seed-soil" hypothesis. These findings illustrate how RGAST can dissect complex tissue microenvironments, offering insights into developmental, homeostatic, and pathological processes.

RGAST’s relational graph architecture addresses a critical gap in ST analysis: the trade-off between spatial fidelity and transcriptional resolution. By jointly modeling spatial neighborhoods and expression profiles, RGAST generates low-dimensional embeddings that outperform state-of-the-art clustering tools (e.g., BayesSpace, STAGATE) in delineating tissue architectures. For instance, in the 10x Visium DLPFC dataset, RGAST improved clustering accuracy (ARI = 0.633) by ∼10% compared to the next-best method, highlighting its utility in identifying spatially coherent domains. Moreover, RGAST’s embeddings facilitated the detection of spatially variable genes (SVGs) with higher Moran’s I score, revealing biologically meaningful patterns (e.g., cortical layer-specific markers) that were less discernible with other methods. Trajectory analysis suggest that RGAST provides a more accurate representation of cell trajectories compared to other methods. Additionally, the 3D RGAST framework, though preliminary, provides a foundation for reconstructing volumetric tissue models from serial sections, mitigating batch effects through cross-section alignment.

By circumventing ligand-receptor database limitations, RGAST provides a systems-level perspective of communication topology, capturing both canonical pathways and novel, context-dependent interactions. This methodological advance positions RGAST as a powerful tool for deconstructing spatial communication networks in complex tissues. RGAST represents a significant step toward comprehensive CCC inference by unifying spatial and transcriptional data into a single, interpretable framework. Its versatility in downstream tasks—from SVG detection to 3D reconstruction—positions it as a powerful tool for unraveling the spatial logic of tissues in health and disease.

## Methods

### Datasets and preprocessing

We applied RGAST to spatial transcriptomics (ST) datasets generated by different platforms, including MERFISH, HDST, 10x Visium, Stereo-seq and seqFISH+. Specifically, the MERFISH dataset was a spatially resolved cell atlas of the mouse hypothalamic preoptic region, profiling ∼1 million cells, each with only 155 genes[15]. We selected six consecutive slices from a single animal sample to conduct our experiments. After removing the putative doublets first, the number of spots ranged from 4,787 to 5,926 for each section. The HDST breast cancer dataset includes 3 tissue sections, which were obtained from a histological grade 3 HER2+ patient [7]. We selected a field of CN21_E2 (spot_x:550∼700, spot_y:550∼650) and removed all cells with unknown cell type or niche annotation. The 10X Visium DLPFC dataset included 12 human DLPFC sections sampled from three individuals. The number of spots ranged from 3,498 to 4,789 for each section, and the original authors manually annotated the areas of DLPFC layers and white matter (WM)[48]. The human intestine dataset was a developmental system dataset from human fetal intestine samples collected at 12 and 19 post-conception weeks (PCW)[59]. We used the A3 sample in our research, which contained only 434 spots. The Stereo-seq mouse olfactory bulb data was binned into a resolution of cellular levels (∼14 μm) and contained 19,109 spots[51]. The MERFISH dataset was a spatially resolved cell atlas of the mouse hypothalamic preoptic region, profiling ∼1 million cells, each with only 155 genes[15]. We selected six consecutive slices from a single animal sample to conduct our experiments. After preprocessing, the number of spots ranged from 4,787 to 5,926 for each section. The mouse cortex seqFISH+ dataset contained mRNAs for 10,000 genes in single cells, with high accuracy and sub-diffraction-limit resolution. This dataset had a spot number of 913[6]. The data sources are summarized in in Supplementary Table S1.

For all datasets, we conducted normalization and scaling, and then ran PCA using the Scanpy package[60]. The PCA representation of each spot was selected as the input of RGAST.

### Construction of relational graph

The spatial neighborhood relationship is established by considering the Euclidean distance between the spatial locations of different spots. In the case of 10x Visium data, we set the graph to include the six nearest neighbors for each spot. For other datasets, we define the adjacency matrix, *A*, such that *A*_*ij*_= 1 if and only if the Euclidean distance between spot *i* and spot *j* is less than a pre-defined hyperparameter, *r*. The expression similarity relationship is constructed by considering the Euclidean distance of the PCA representation of the gene expression spectra of different spots.

### Relational graph attention auto-encoder

The relational graph attention auto-encoder consists of an encoder of two relational graph attention layer and a linear layer decoder.

#### Encoder

The input of the encoder in our architecture is a graph with *R* = 2 relation types and *N* nodes (spots). The *i*_*th*_ node is represented by a feature vector of the PCA dimensional reduction representation of gene expressions ***x***_*i*_. Query and key representations for the *l*_*th*_ encoder layer are computed for each relation type with the help of both query and key kernels, i.e.

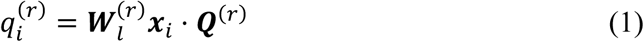

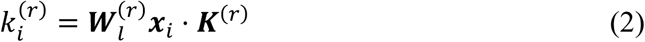

where ***W***_*l*_ is the trainable weight matrix of layer *l*. Then additive attention[61] is applied to compute attention logits 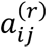:

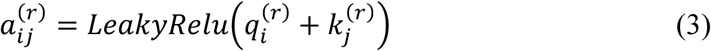

Then the attention coefficients for each relation type are then obtained via the across-relation attention mechanism:

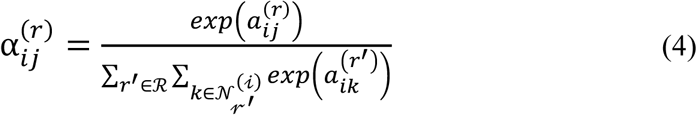

where ℛ denotes the set of relations, i.e. edge types. 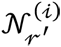 denotes the set of spots connected to node *i* under relation *r*′. To avoid overfitting, we employed the dropout strategy on the normalized attention coefficients with dropout rate=0.3. That is, 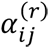 will be set as 0 with a probability of 0.3.

Then spot *i* collectively aggregating information from spots connected to it with the neighborhood aggregation step to get the output of layer *l*. We denote the intermediate representation of 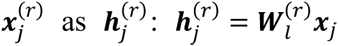. To enhance the discriminative power of the RGAST layer, we further implement the additive cardinality preservation mechanism[62]:

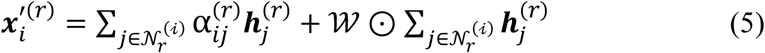

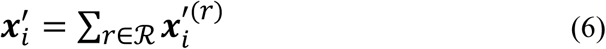

where 𝒲 is a non-zero vector ∈ *R*^*n*^, n is the output dimension of layer *l*, ⊙ denotes the elementwise multiplication. The output of encoder is considered as the final spot embedding.

#### Decoder

The decoder reverses the latent embedding back into the original PCA representation of the expression profile. The one-layer linear decoder treats the output of the encoder (denoted by ***h***_*i*_) as its input and computes the reconstructed result. Specifically, the decoder computes the reconstructed result as follows:

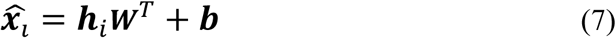

Where ***W*** and ***b*** are learnable weight matrix and bias vector.

#### Loss function

The objective of RGAST is to minimize the reconstruction loss of the original PCA profiles as follows:

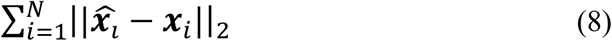

In all experiments we used Adam optimizer with learning rate=0.001 and weight decay=0.0001.

### Deep embedding clustering (optional)

To enhance the compactness of the RGAST embedding, we use an unsupervised deep embedded clustering (DEC) method to iteratively group cells into different clusters[14]. To initialize the cluster centers, we first apply a clustering method to the learned latent representations (such as Louvain, Leiden, Kmeans, etc.). With this initialization, DEC improves the clustering using an unsupervised iterative method of two steps. In the first step, DEC calculates a soft assignment *q*_*ij*_ of the latent point *h*_*i*_ to the cluster center mui using the Student’s t-distribution:

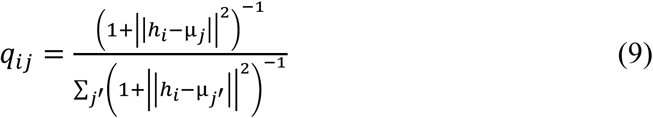

In the second step, we iteratively refine the clusters by learning from their high confidence assignments with the help of an auxiliary target distribution *p* based on *q*_*ij*_:

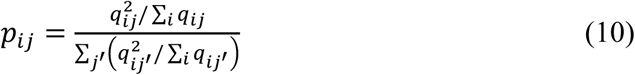

Based on the soft assignment *q*_*ij*_ and auxiliary target distribution *p*_*ij*_, an objective function is defined using the KL divergence:

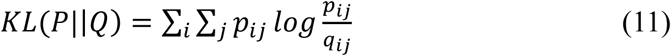

Then the overall loss function is the sum of (8) and (11). The RGAST parameters and cluster centers are then simultaneously optimized using Adam optimizer.

### Scalable training of RGAST

Training RGAST becomes challenging when the number of spots is large, as mini-batch training is not well-defined for graph neuron networks. Here we adopted the recently developed Divide-Iterate-Conquer (DIC) strategy for large scale spatial transcriptomics, which training on subgraphs and inferring on the whole slice [56]. In our Stereo-seq mouse olfactory bulb dataset, we divided the graph into 2×2 subgraphs. The dividing point along the x-axis is determined by the median of spot locations, which is the same for the y-axis.

### Multi-scale cell-cell communications analysis

#### Single-cell level

Following model training, each relational graph attention convolution (RGATConv) layer generates edge-specific attention scores within the original heterogeneous graph. For our analysis, we specifically utilized the attention scores from the final RGATConv layer to reconstruct the cell-cell communication (CCC) network. To construct a biologically meaningful CCC network, we implemented an attention score thresholding approach. For each individual cell/spot, we: 1. Sorted all connecting edges in descending order based on their attention scores; 2. Computed the cumulative sum of attention scores; 3. Retained edges only until the cumulative sum exceeded a predetermined threshold. This selective filtering process effectively preserves the most relevant interactions while eliminating weaker, potentially noisy connections. The resulting pruned graph represents a high-confidence, single-cell resolution CCC network that captures the most significant intercellular communication events.

#### Niche level

To characterize niche-level signaling patterns, we derived spatial vector fields from our reconstructed single-cell CCC network. Given the reconstructed attention score matrix ***S*** ∈ 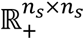, where ***S***_*ij*_ represents the directed signaling strength from spot *i* to spot *j*, we computed two distinct vector fields: Signal-sending vector field (***V***^*s*^) and Signal-receiving vector field ( ***V***^*r*^). For each spot *i*, we define: 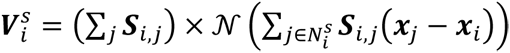, 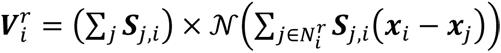, where 𝒩(***x***) = ***x***‖***x*** denotes the normalization operator, 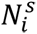 represents the index set of top k signal-sending neighbors for spot *i* (highest values in row *i* of ***S***), 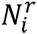 represents the index set of top k signal-receiving neighbors for spot *i* (highest values in column *i* of ***S***). ***x***_*i*_ ∈ ℝ^2^ denotes the spatial coordinates of spot *i*.

#### Cell type level

To derive cell type-level cell-cell communication (CCC) metrics, we implemented a normalized counting approach that accounts for cellular composition biases. For each pairwise interaction between cell types A and B, we: 1. Enumerated all intercellular connections (edges) between the two cell types; 2. Normalized the raw connection count by the product of their respective cell population sizes ( *N*_*A*_ × *N*_*B*_); 3. Defined this normalized value as the communication score (*CS*_*AB*_): *CS*_*AB*_ = (Number of connections between A and B) / (*N*_*A*_ × *N*_*B*_). The communication score mitigates the confounding effects of unequal cell type abundances and provides an unbiased metric for comparing communication strengths across different cell type pairs.

### Benchmarking different CCC methods

In real data, single-cell level CCC network ground truth does not exist. Since all the compared methods are capable of generating single-cell-level cell-cell communication (CCC) matrices, we constructed a proxy CCC matrix as a plausible ground truth through a voting mechanism: for each cell pair, we determined the presence of an edge if at least half of the methods agreed on its existence. Although straightforward, this meta-learning-inspired approach yields high-confidence results. Using this high-confidence CCC matrix as the ground truth, we computed standard binary classification metrics (e.g., precision, recall, F1-score, and accuracy) to evaluate model’s performance.

### Spatial domain detection / clustering analysis

#### Leiden algorithm

We used the Scanpy package to compute the Leiden clustering from the latent embeddings generated from different representation learning methods. Briefly, we used latent embeddings to compute neighborhood graphs with number of neighbors as the recommended default parameter and Euclidean distance as the distance measure. To set the maximum number of unique Leiden clusters, grid-searching on the Leiden clustering resolutions between 0.0 and 2.5 was performed at intervals of 0.02.

#### Evaluation metrices

##### Adjusted Rand index (ARI)

The Rand Index computes a similarity measure between two clusterings by considering all pairs of samples and counting pairs that are assigned in the same or different clusters in the predicted and true clusterings. The raw RI score is then “adjusted for chance” into the ARI score using the following scheme:

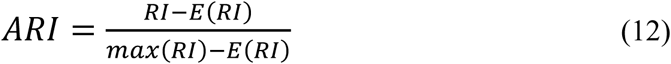

To calculate this value, first calculate the contingency table like that:

| | $Y_1$ | $Y_2$ | ... | $Y_s$ | Sums |
| --- | --- | --- | --- | --- | --- |
| $X_1$ | $n_{11}$ | $n_{12}$ | ... | $n_{1s}$ | $a_1$ |
| $X_2$ | $n_{21}$ | $n_{22}$ | ⋯ | $n_{2s}$ | $a_2$ |
| ⋮ | ⋮ | ⋮ | $\ddots$ | ⋮ | ⋮ |
| $X_r$ | $n_{r1}$ | $n_{r1}$ | ⋯ | $n_{rs}$ | $a_r$ |
| Sums | $b_1$ | $b_2$ | ⋯ | $b_s$ | |

each value in the table represents the number of data point located in both cluster (Y) and true class (X), and then calculate the ARI value through this table:

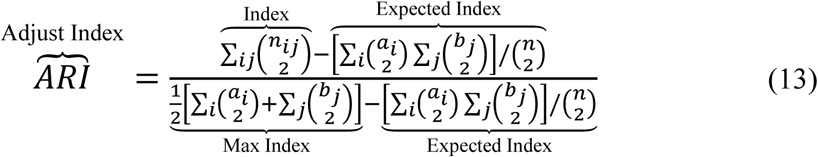

The adjusted Rand index is thus ensured to have a value close to 0.0 for random labeling independently of the number of clusters and samples and exactly 1.0 when the clusterings are identical (up to a permutation). The adjusted Rand index is bounded below by −0.5 for especially discordant clusterings.

##### Silhouettes Coefficient (SC)

SC is a metric used to evaluate the clustering effectiveness on dataset without ground truth label[63]. *a*_*i*_ is denoted as tightness, which represents the average distance between sample *i* and other samples within same cluster. The average distance between one sample and other samples from different cluster is defined as separation. *b*_*i*_ = *min*[*b*_*i*1_, *b*_*i*2_,…, *b*_*ik*_], which is the separation of sample *i*. Then the Silhouette Coefficient of sample *i* is defined as:

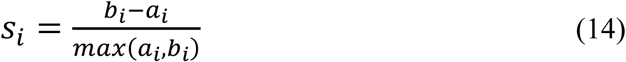

The SC score is the average of *s*_*i*_ ranging from −1 to 1 and higher score means higher performance.

### Spatially variable genes detection

#### Detection of SVGs

In this study, we employ the same detection method as that used in SpaGCN to identify spatially variable genes (SVGs) enriched in each spatial domain generated by different methods. We recognize that some genes may be expressed in multiple but disconnected domains. Therefore, instead of conducting differential expression (DE) analysis using spots from a target domain versus all other spots, we first select spots to form a neighboring set around the target domain. Our aim is to detect genes that are highly expressed in the target domain but not expressed or expressed at low levels in neighboring spots. To achieve this, we draw a circle with a predetermined radius around each spot in the target domain, and spots from non-target domains within the circle are considered its neighbors. The radius is set such that each spot in the target domain has an average of approximately ten neighbors. Next, we collect all the neighbors of spots in the target domain to form a neighboring set. For each non-target domain, if more than 50% of its spots are in the neighboring set, we select it as a neighboring domain. This criterion is in place to avoid cases where a domain is selected as a neighboring domain, but only a small proportion of its spots are adjacent to the target domain. Once neighboring domains are determined, we perform DE analysis between spots in the target domain and the neighboring domain(s) using the Wilcoxon rank-sum test. Genes with a false discovery rate (FDR)-adjusted P-value < 0.05 are selected as SVGs. To ensure only genes with enriched expression patterns in the target domain are selected, we require a gene to meet the following three criteria: (1) the percentage of spots expressing the gene in the target domain, that is, in-fraction, is > 80%; (2) for each neighboring domain, the ratio of the percentages of spots expressing the gene in the target domain and the neighboring domain(s), that is, in/out fraction ratio, is > 1; and (3) the expression fold change between the target and neighboring domain(s) is > 1.5.

#### Evaluation metrices

Gene expressions at different locations may not be independent. For example, the expression levels of a gene at nearby locations may be closer in value than expression levels at locations that are farther apart. This phenomenon is called spatial autocorrelation, which measures the correlation of a variable with itself through space. To evaluate whether the detected SVGs exhibit an organized spatial expression pattern, we used Moran’s I and Geary’s C, two commonly used statistics to quantify the degree of spatial autocorrelation of gene expression.

##### Moran’s I statistic

Moran’s I metric is a correlation coefficient that measures the overall spatial autocorrelation of a dataset[64]. Essentially, it quantifies how similar one spot is to other spots surrounding it for a given gene. If the spots are attracted or repelled by each other, it suggests that they are not independent, and the presence of autocorrelation indicates a spatial pattern of gene expression. The Moran’s I value ranges from −1 to 1, where a value close to 1 indicates a clear spatial pattern, a value close to 0 indicates random spatial expression, and a value close to −1 indicates a chessboard-like pattern. To evaluate the spatial variability of a given gene, we calculate the Moran’s I using the following formula:

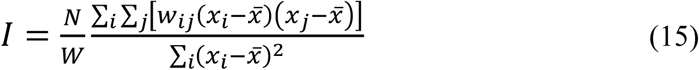

where *x*_*i*_ and *x*_*j*_ are the gene expression of spots *i* and *j*, *x̄* is the mean expression of the gene, *N* is the total number of spots, *w*_*ij*_ is the spatial weight between spots *i* and *j* calculated using the 2D spatial coordinates of the spots, and *W* is the sum of *w*_*ij*_. To select the k nearest neighbors for each spot, we use spatial coordinates. The Moran’s I statistic is robust to the choice of k, and we set it to 5 in our analysis. We assign *w*_*ij*_=1 if spot *j* is in the nearest neighbors of spot *i*, and *w*_*ij*_=0 otherwise.

##### Geary’s C statistic

Geary’s C statistic is another measure of spatial autocorrelation, which is a statistical method used to examine the degree to which neighboring observations in a dataset are similar to each other for a given gene[65]. In other words, it measures the extent to which spots that are close to each other in space are more similar or dissimilar than those that are farther apart with respect to the gene expression. Geary’s C is calculated using the following formula:

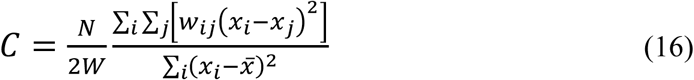

The value of Geary’s C ranges from 0 to 2, with a value of 1 indicating no spatial autocorrelation, values less than 1 indicating positive spatial autocorrelation (i.e., neighboring expressions are more similar than expected by chance), and values greater than 1 indicating negative spatial autocorrelation (i.e., neighboring expressions are more dissimilar than expected by chance).

### Trajectory and pseudotime inference

#### PAGA graph and UMAPs

We also used the Scanpy package to compute the partition-based graph abstraction (PAGA), and uniform manifold approximation and projection (UMAP) from embeddings generated from different methods. After using latent embeddings to compute neighborhood graphs, UMAP algorithm was conducted based on the neighborhood graphs. Finally, PAGA was applied to quantify the connectivity between different layers based on the UMAP representations.

#### Monocle3

For SEDR, STAGATE, conST and RGAST, we first extracted the low-dimensional embedding and then used the embedding to replace the default PCs used in Monocle3 package. We then ran Monocle3 on UMAPs generated by the latent embedding using the recommended parameters and randomly set a spot in white matter (WM) as the starting point to generate the pseudo-time.

### 3D RGAST model

All current ST technologies capture gene expression patterns within the context of 2D tissue sections, which limits the accurate representation of 3D spatial information in real-world samples. A conventional approach to address this limitation is by reconstructing gene expressions in 3D space through the stacking of consecutive 2D sections[3, 66]. However, the presence of batch effects between sections poses a challenge in extracting meaningful 3D spatial patterns. Here, we proposed a 3D RGAST model that incorporates both the 2D relationships within each section and the spatial neighborhood relationships between adjacent sections to mitigate the impact of batch effects. If the presence of minimal batch effects between different sections is confirmed, gene expression similarity can also be applied across sections. Specifically, the spatial neighborhoods between adjacent sections are constructed based on aligned coordinates and a pre-defined radius. The underlying principle behind the use of 3D spatial neighborhoods lies in the assumption that biological differences between consecutive sections should demonstrate continuity. By enhancing the similarity between adjacent sections, we aim to eliminate discontinuous independent technical noises.

## Declarations

### Data and code availability

All data used in this research can be found in **Supplementary Table S1**. Our RGAST method is available as a Python package on PyPI at https://pypi.org/project/RGAST, free for academic use, and the source code is openly available from our GitHub repository at https://github.com/GYQ-form/RGAST.

### Authors’ contributions

YG performed the research, analyzed data and wrote the original manuscript, ZY supervised the research. YG and YZ discussed and revised the manuscript.

## Acknowledgements

We are thankful for Xin Yuan for providing practical suggesting for the research. The computations in this paper were run on the π 2.0 cluster supported by the Center for High Performance Computing at Shanghai Jiao Tong University.

