## Supplementary Information for "RGAST: A Relational Graph Attention Network for Multi-Scale Cell-Cell Communication Inference from Spatial Transcriptomics"

### Supplementary Tables

#### Table S1. Detailed description of the datasets used in the paper

| Datasets | Description | Source |
|---|---|---|
| Mouse hypothalamic preoptic | six consecutive slices from a single mouse's preoptic region of hypothalamus profiled by MERFISH | https://datadryad.org/stash/dataset/doi:10.5061/dryad.8t8s248 |
| Human breast cancer | HDST data obtained from a histological grade 3 HER2+ patient | https://portals.broadinstitute.org/single_cell/study/SCP420 |
| Human DLPFC | 12 human DLPFC sections sampled from three individuals profiled by 10X Visum | http://spatial.libd.org/spatialLIBD/ |
| Mouse olfactory bulb | Data from mouse olfactory bulb tissues profiled by Stereo-seq | https://github.com/JinmiaoChenLab/SEDR_analyses |
| Mouse cortex | seqFISH+ dataset contained mRNAs for 10,000 genes in single cells, with high accuracy and sub-diffraction-limit resolution | https://www.spatialomics.org/SpatialDB/seqfifish_30911168_browse.php |

### Supplementary Figures

#### Fig S1-S11. clustering results for the remaining 10X Visium DLPFC slices

151507

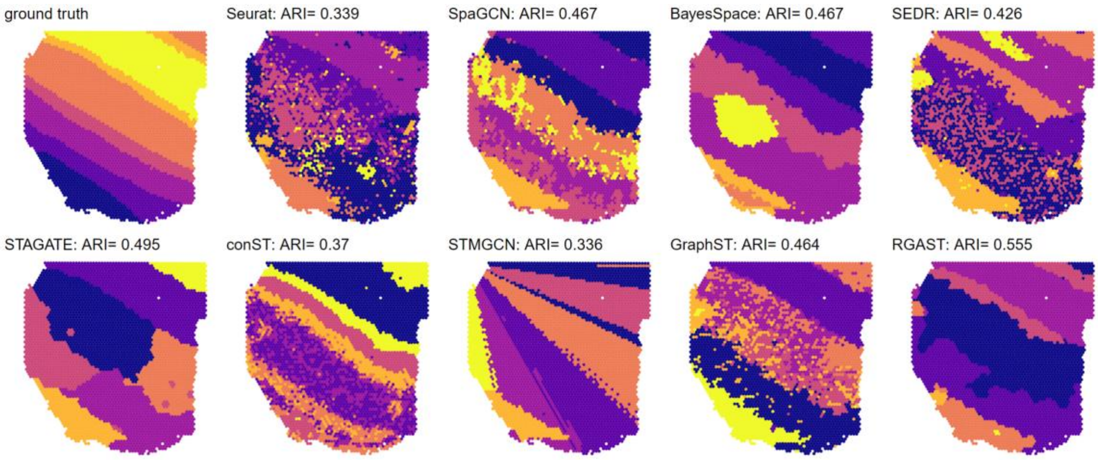


151508

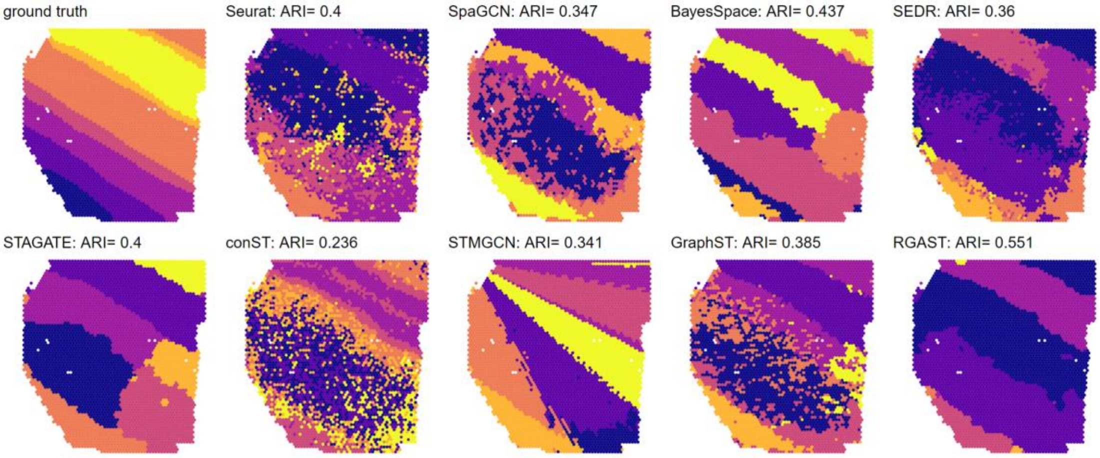


151509

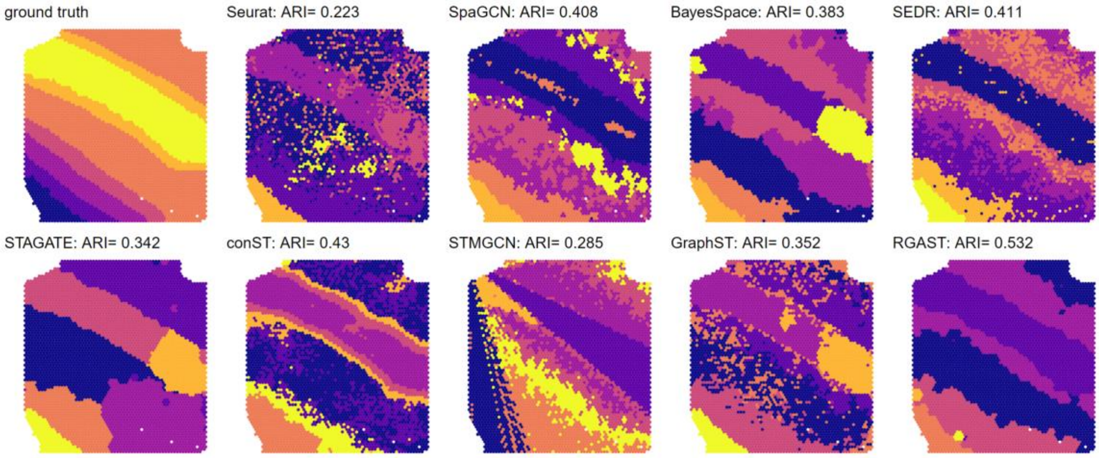

151510

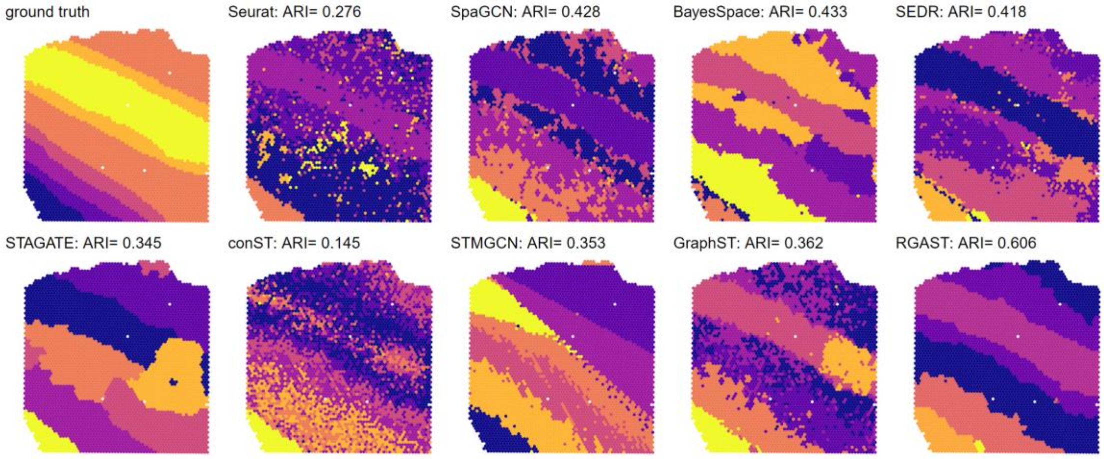


151669

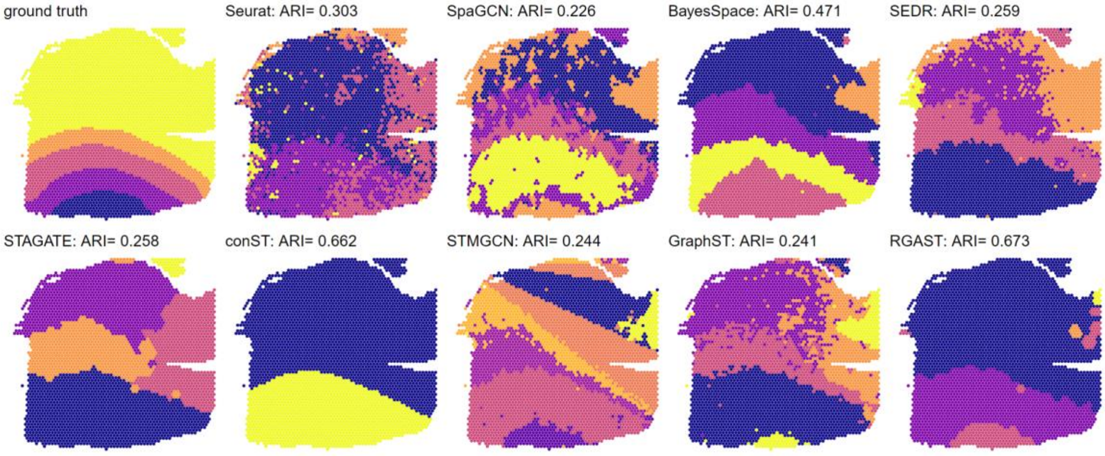


151670

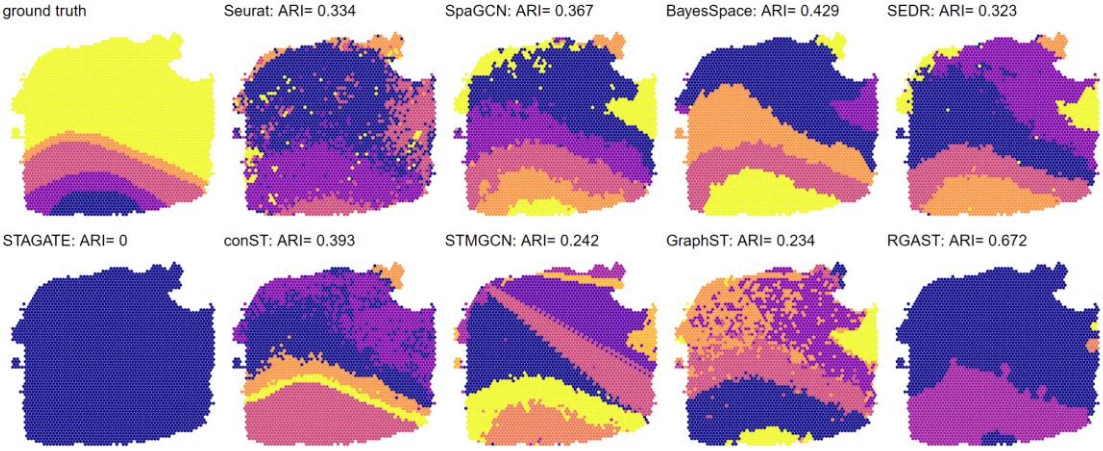

151671

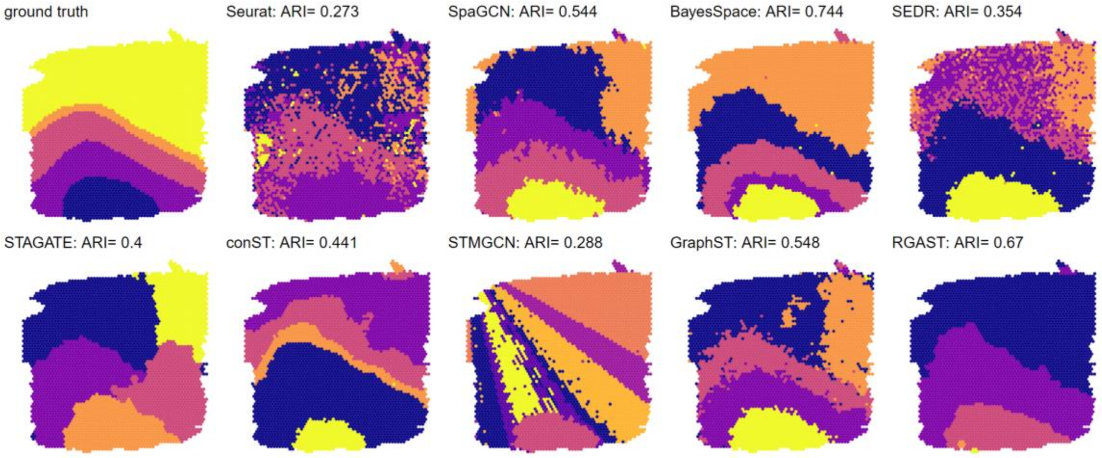


151672

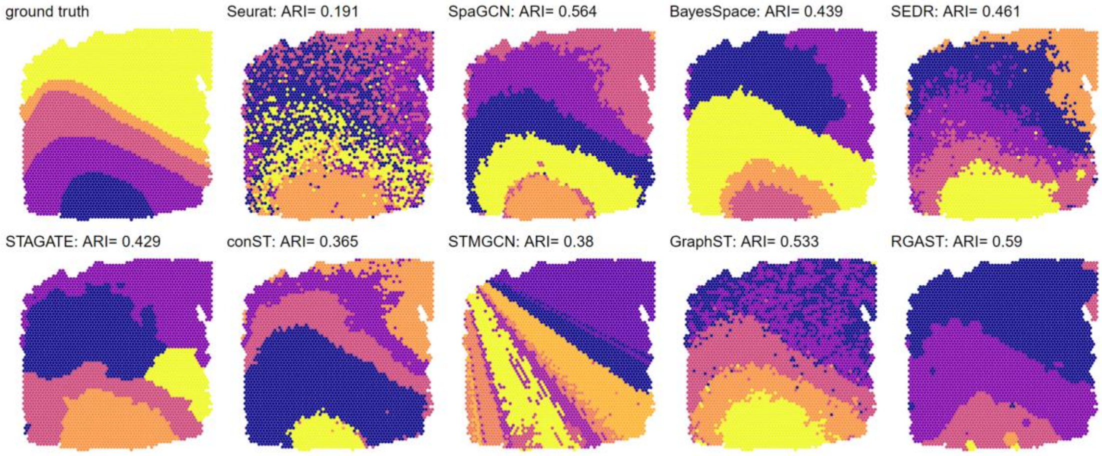


151673

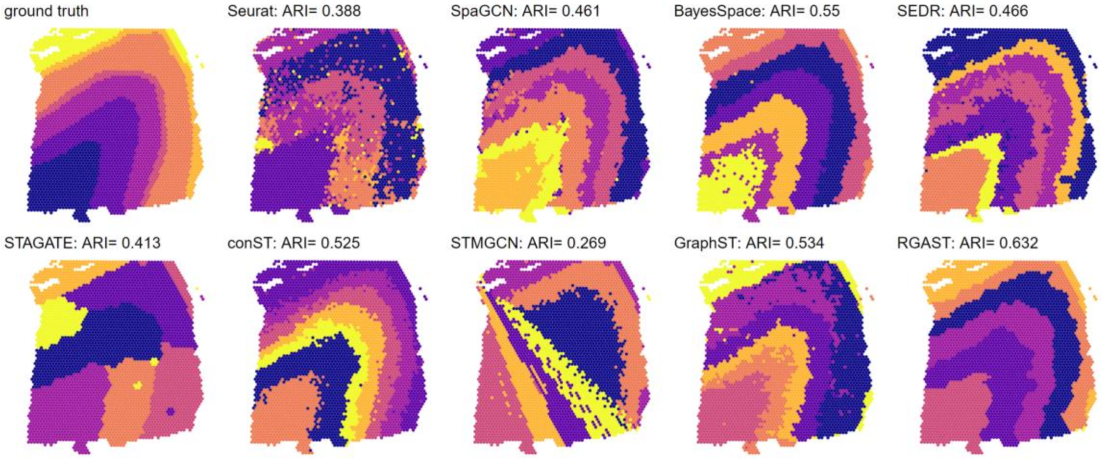

151674

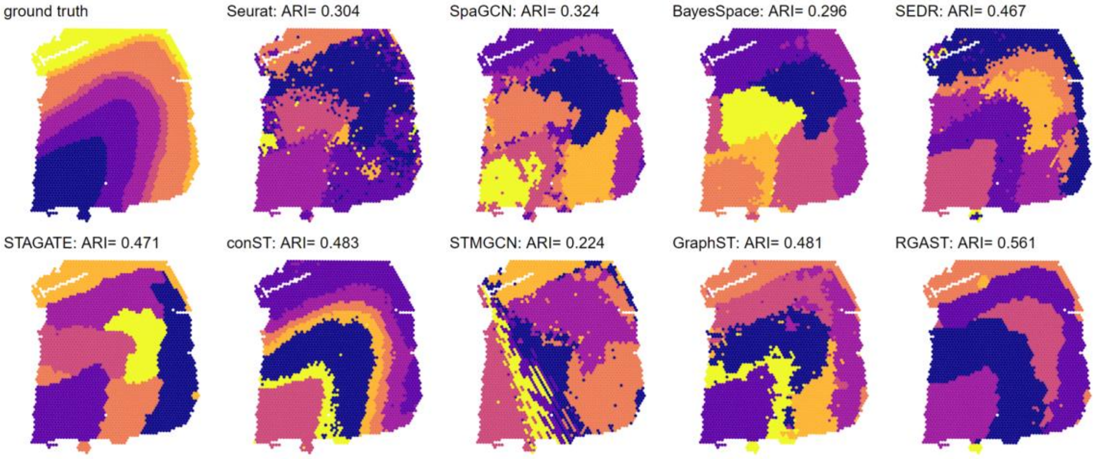


151676

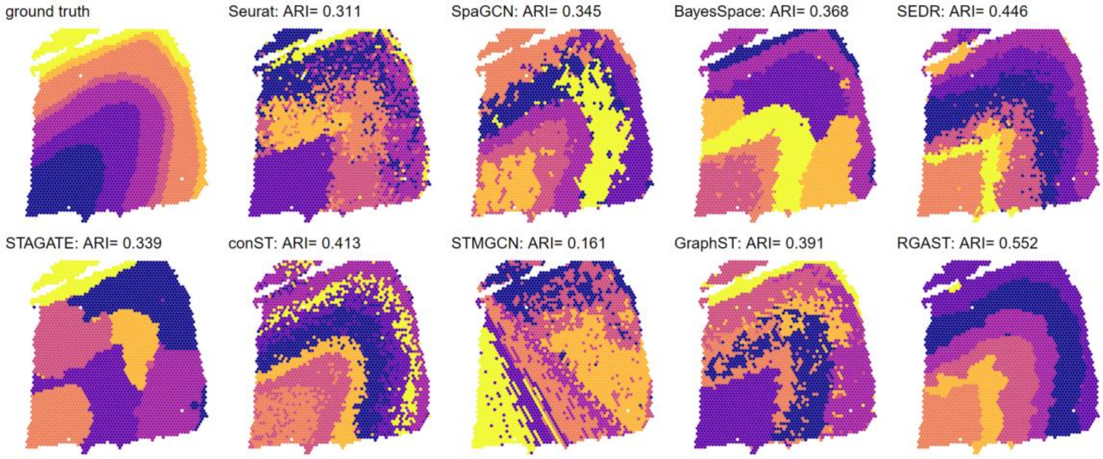

**Fig S12. Spatial domains and corresponding marker genes identified by RGAST in Stereo-seq mouse olfactory bulb tissue**

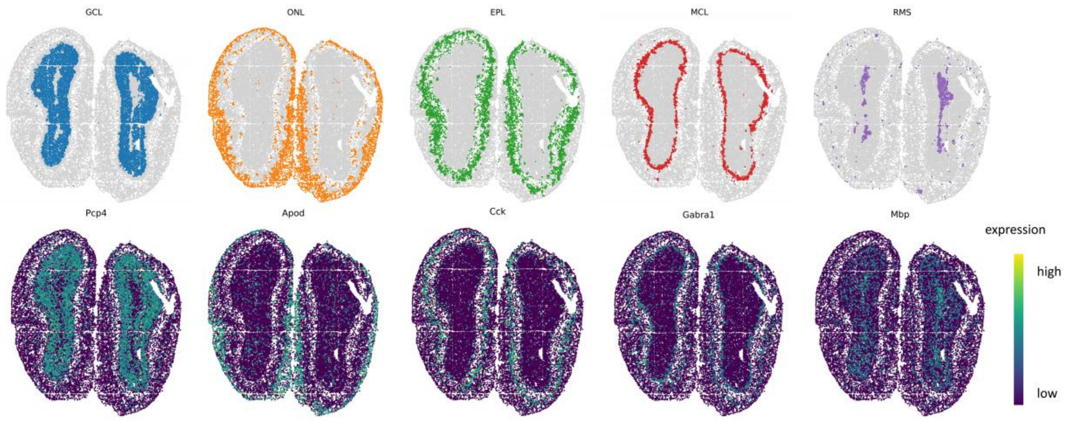

**Fig S13-S17. clustering results for the remaining MERFISH mouse hypothalamic preoptic region slices**

Bergman 0.16

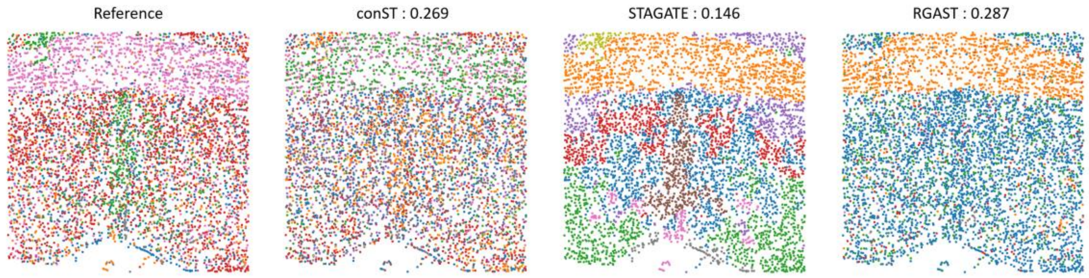


Bergman 0.06

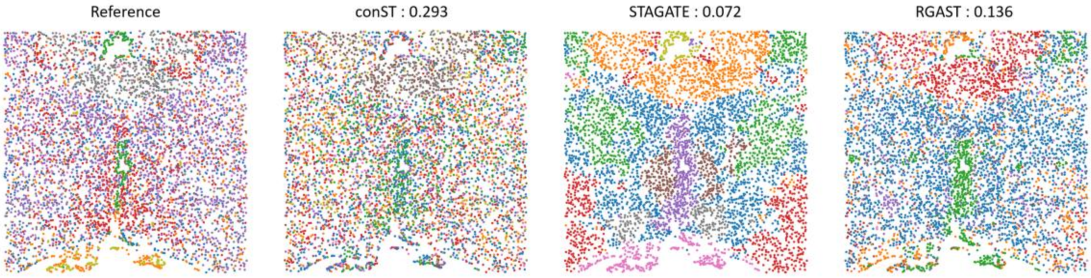


Bergman -0.04

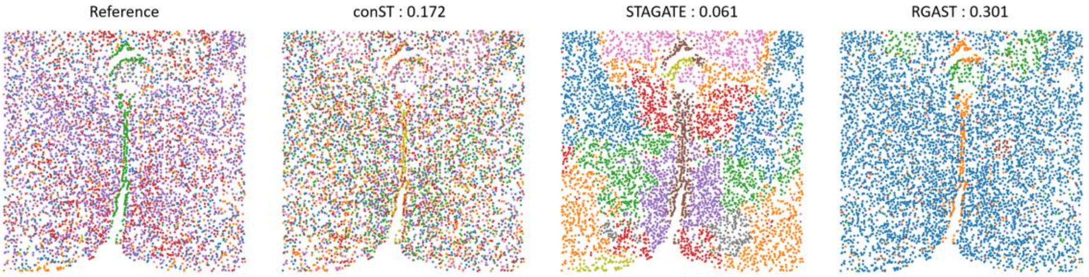


Bergman -0.14

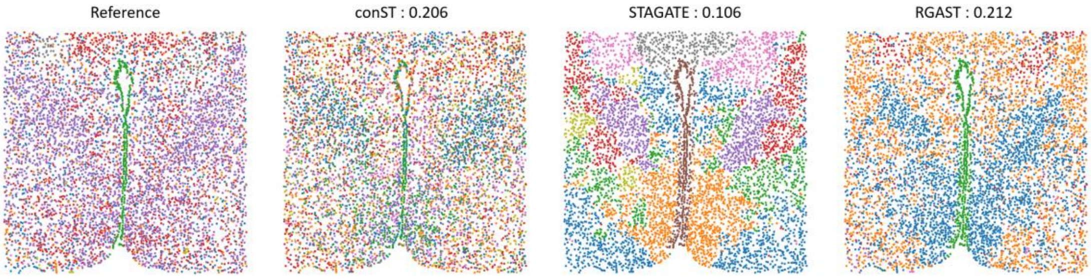

Bergman -0.24

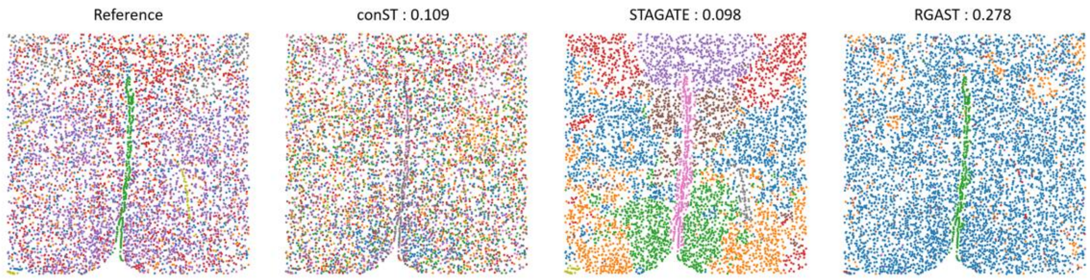

**Fig S18. Heatmap of gene expression in the merged MERFISH data**

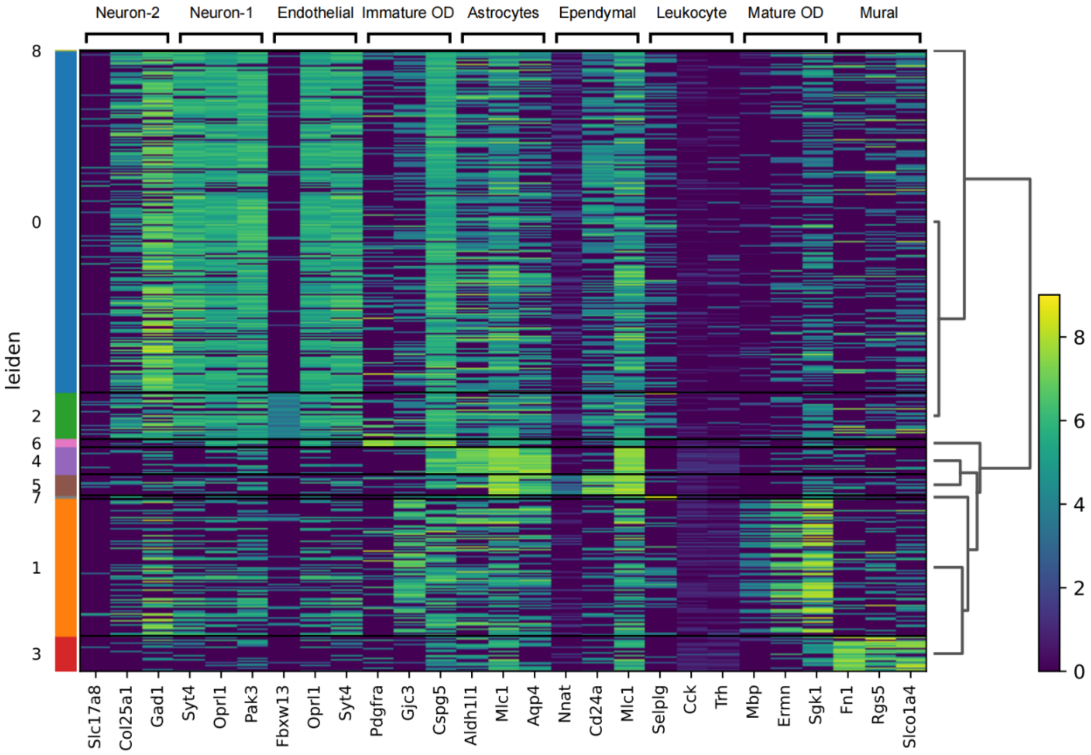

### Fig S19. 3D Spatial domain of MERFISH data identified by RGAST

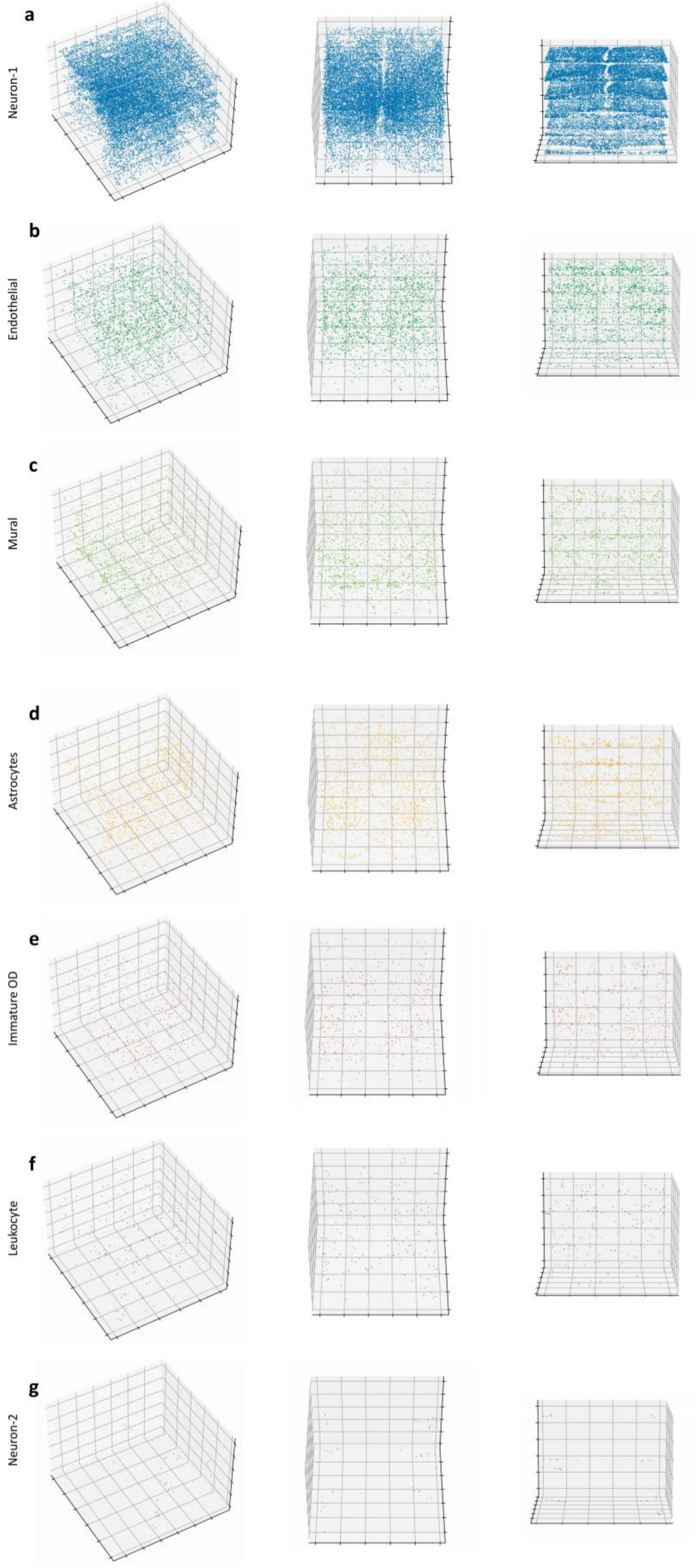
